# NOLC1 Suppresses Immuno-chemotherapy by Inhibiting p53-mediated Ferroptosis in Gastric Cancer

**DOI:** 10.1101/2024.11.05.622078

**Authors:** Shengsheng Zhao, Ji Lin, Bingzi Zhu, Yin Jin, Qiantong Dong, Xiaojiao Ruan, Dan Jin, Yongdong Yi, Binglong Bai, Hongzheng Li, Danna Liang, Jianhua Lu, Letian Meng, Xiang Wang, Yuekai Cui, Yuyang Gu, Xian Shen, Xufeng Lu, Shangrui Rao, Weijian Sun

## Abstract

Gastric cancer (GC) is one of the most malignant cancers, and cisplatin (Cis)-based chemotherapy remains the main clinical treatment for GC. However, Cis resistance often occurs, largely limiting its therapeutic efficacy in tumors. Therefore, a better understanding of the drug resistance mechanism could reveal new approaches for improving GC treatment efficacy. Here, we define the integrative role of nucleolar and coiled-body phosphoprotein 1 (NOLC1), a molecular chaperone that is significantly upregulated in GC tissues and Cis-resistant GC cells. Knocking down NOLC1 increased GC sensitivity to Cis by regulating ferroptosis. Mechanistically, NOLC1 binds to the p53 DNA binding domain (DBD), decreasing p53 nuclear accumulation stimulated by Cis and suppressing p53 transcriptional functions. Then, the p53-mediated ferroptosis is suppressed. Furthermore, the silence of NOLC1 promoted ferroptosis-induced immunogenic cell death (ICD) and reprogrammed the immunosuppressive tumor microenvironment, thereby increasing sensitivity to anti-programmed cell death-1 (PD-1) therapy plus Cis. The combination of anti-PD-1 plus Cis effectively inhibited GC growth without significant side effects. In summary, our findings reveal that targeting NOLC1 may be a novel therapeutic strategy for GC and may increase the efficacy of chemotherapy combined with immune checkpoint inhibitor (ICI) therapy.

## Introduction

Gastric cancer (GC) is the fifth most diagnosed cancer worldwide and the fifth most common cause of cancer-related death^1^. Since GC is usually diagnosed at an advanced stage, it often has a poor prognosis^2,3^. Cisplatin (Cis)-based chemotherapy remains the primary treatment for advanced GC^4^. Chemotherapy not only has a direct cytotoxic effect but also contributes to the immunogenic cell death (ICD) and improves the efficacy of immunotherapy. Thus, immunotherapy, represented by anti-programmed cell death-1 (PD-1) therapy, plus chemotherapy, has become the standard for first-line treatment^2,5^. However, the efficacy of chemotherapy is still restricted by chemoresistance, which is caused by gene mutations, gene inactivation, epigenetic modifications, signaling pathway changes, or cell metabolism disorders^6^; thus, the efficacy of immune-chemotherapy is still limited in some patients^7^. Therefore, exploring novel targets of chemoresistance in GC to increase combination effectiveness is highly important for improving tumor prognosis and patient survival.

Ferroptosis, an iron-dependent form of novel regulated cell death (RCD), is regulated by various pathways, such as mitochondrial activity, oxidative stress, iron handling, and metabolism^8,9^. Recently, studies have shown that activation of ferroptosis in GC is a promising strategy for overcoming GC resistance. For example, increased ATF2 expression can alleviate Cis resistance by regulating ferroptosis in GC^10^. Additionally, GC cells can overexpress many negative ferroptosis regulators, such as miR-522 and ACTL6A, to promote chemoresistance^11,12^. Moreover, ferroptotic cells can activate the immune response and reprogram the immunosuppressive tumor microenvironment (TME) via releasing various damage-associated molecular patterns (DAMPs), subsequently increasing the efficacy of immunotherapy^13,14^. Thus, ferroptosis may serve as a new strategy for increasing the efficacy of immunotherapy combined with Cis. It is well established that p53, a classic tumor suppressor protein, can regulate ferroptosis via multiple mechanisms^15,16^. As a traditional transcription factor (TF), p53 transcriptionally regulates several metabolic or ferroptotic targets, such as TIGAR, GLS2, and ALOX12, to mediate ferroptosis^17–19^. However, inactivation of p53 tumor suppressor function is very common and plays an important role in the progression of most cancers^20^. Among patients with GC, those with activated p53 status have longer overall survival than those with p53 loss of function^21^. Recently, studies have shown that activating p53 can increase chemotherapy efficacy by promoting ferroptosis^22,23^. Therefore, activating p53 function in GC is a highly desired feature of ferroptosis-based antitumor therapy.

Nucleolar and coiled-body phosphoprotein 1 (NOLC1), also called NOPP140, was originally identified as a nuclear localization signal binding protein (NLS) and functions as a molecular chaperone that shuttle between the cytoplasm and nuclear^24,25^. NOLC1 is up-regulated in most cancers and promotes non-small cell lung cancer (NSCLC) resistant to multiple drugs^26,27^. Moreover, NOLC1 can interact with the MDM2 promoter at the p53 binding site, and NOLC1 was also identified mediate by p53 via an RNAi-mediated synthetic interaction screen^28,29^. However, the specific expression level and functions of NOLC1 in GC remains unclear.

Here, we found that NOLC1 is up-regulated in GC tumors and GC resistant cells. High expression of NOLC1 promotes GC Cis resistance by suppressing ferroptosis. Further studies revealed that NOLC1 interacts with the p53 DNA Binding Domain (DBD) and suppress p53 nuclear accumulation, thus inhibiting p53 transcription-mediated ferroptosis. Eventually, the knockdown of NOLC1 activated the ferroptosis-mediated ICD and significantly improved the efficacy of low-dose anti-PD-1 plus Cis without side effects. Collectively, this study identified a novel network mediated by NOLC1 that regulates p53-mediated ferroptosis in GC and identified a new target for GC immunotherapy in combination with Cis.

## Results

### High expression of NOLC1 is associated with poor clinical outcomes in GC

Recent studies have demonstrated that NOLC1 is up-regulated in most cancers^27^. However, the expression of NOLC1 in GC is still unknown. To address this gap in knowledge, we collected GC patients’ tumor and near-tumor tissues to measure NOLC1 expression level. As is shown in **Fig. 1A and Supplementary Fig. S1 A,** immunohistochemical (IHC) staining demonstrated that NOLC1 protein level was much higher in GC tissues than in near-tumor tissues. Compared with that of normal tissues, the average optical density (AOD) score was notably greater in GC tissues (**Fig. 1B**). Consistently, as is shown in **Fig. 1C-D**, western blotting (WB) results also revealed the same trend: NOLC1 was highly expressed in GC tissues. Analysis of the TCGA database also revealed that NOLC1 is up-regulated in GC tissues (**Fig. 1E-F**). We next used the Kaplan□Meier plotter database to assess whether high expression of NOLC1 influenced GC patient outcomes. Higher NOLC1 expression level was related to shorter overall survival (OS), time to first progression (FP), and postprogression survival (PPS) in patients with GC (**Fig. 1G-I**). Taken together, all these results suggest that NOLC1 protein levels are increased in GC and that NOLC1 may promote the progression of GC.

**Fig. 1.**
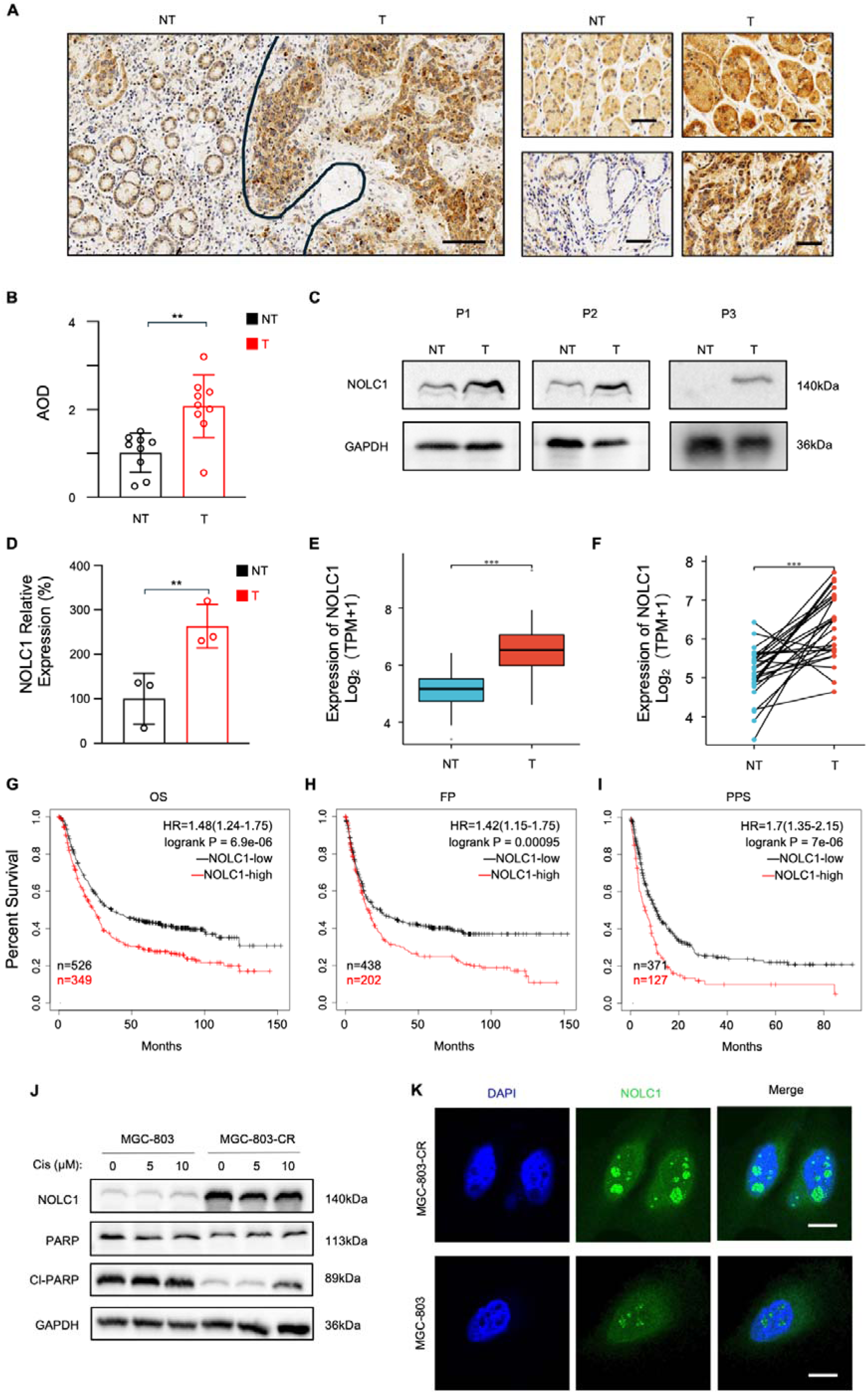
NOLC1 was up-regulated in GC and a Cis-resistant GC cell line. A, **B** Immunohistochemistry (IHC) results of NOLC1 expression in gastric tumor (T) tissues and near tumor (NT) tissues. (A) Representative images (*n* = 3); scale bar = 50 μm. (B) Average density (AOD) of NOLC1 according to the IHC images (n = 9). **C, D** Western blotting (WB) results of NOLC1 protein levels in gastric tumor (T) tissues and near-tumor (NT) tissues. (C) Representative images. (D) Relative expression of NOLC1 (*n* = 3). **E, F** Gene expression level of NOLC1 in TCGA GC tumor and matched TCGA normal stomach tissues. (E) Unpaired sample, (F) paired sample. **G-I** High NOLC1 expression predicts poor survival in GC patients. (G) Overall survival (OS); patients were divided into a NOLC1-high group (*n* = 349) and a NOLC1-low group (*n* = 526). (H) Time to first progression (FP); patients were divided into a NOLC1-high group (*n* = 202) and a NOLC1-low group (*n* = 438). (I) Postprogression survival (PPS); patients were divided into a NOLC1-high group (*n* = 127) and a NOLC1-low group (*n* = 371). **J** WB results of NOLC1, PARP, and cleaved PARP in MGC-803 and MGC-803-CR cell lines. **K** IF images of NOLC1 in MGC-803 and MGC-803-CR cell lines; scale bar = 15 μm. The data are presented as the means ± SDs. **p < 0.01; ***p < 0.001.

Since NOLC1 is highly expressed and promotes resistance to multiple drugs in NSCLC^26^, we further detected whether NOLC1 is associated with GC resistance. To investigate the expression of NOLC1 in Cis-resistant GC, a Cis-resistant MGC-803 (MGC-803-CR) was established. CCK-8, colony formation, and annexin V-APC/7-AAD apoptosis assays revealed that MGC-803-CR cells were less sensitive to Cis than MGC-803 cells (**Supplementary Fig. S2 A-E)**. Next, the expression of NOLC1 in MGC-803-CR cells was detected. WB showed that NOLC1 is highly expressed in MGC-803-CR cells (**Fig. 1J, Supplementary Fig. S3A**). The cleaved PARP protein level was significantly lower than that in MGC-803 cells (**Supplementary Fig. S3B**), further supporting that MGC-803-CR cells are resistant to Cis. Moreover, immunofluorescence analysis revealed that NOLC1 was upregulated in both the cytoplasm and nucleolus (**Fig. 1K, Supplementary Fig. S3C**). These data indicate that NOLC1 is highly expressed in the Cis-resistance cell line. In addition, we used mRNA-seq to analyze the specific biological pathway alterations in MGC-803-CR cells. As shown in **Supplementary Fig. S3 D, E**, the expression level of components of the Hippo signaling pathway, PI3K-AKT pathway, p53 signaling pathway, etc., significantly differed between MGC-803-CR cells and MGC-803 cells.

### NOLC1 knockdown increases the sensitivity of GC to Cis in *vitro* and in *vivo*

To investigate the functions of NOLC1 in Cis-resistance GC, lentiviral shRNA constructs (shNOLC#1, #2, or #3) used to knock down endogenous NOLC1 gene expression in GC cell lines, MGC-803 and MKN-45 cells, respectively (**Supplementary Fig. S4 A-D**). CCK-8 assay revealed that NOLC1 silencing significantly decreased the viability of MGC-803 and MKN-45 cells upon Cis treatment (**Fig. 2A**). Meanwhile, overexpression (OE) of NOLC1 promoted Cis resistance in MGC-803 cells (Supplementary Fig. S5A). The colony formation assay results also revealed that knockdown NOLC1 could significantly decreased the numbers of colonies after Cis treatment (**Fig. 2B-C**). Moreover, the annexin V-APC/7-AAD apoptosis assay revealed that Cis induced cell death to a greater degree in the NOLC1 knockdown group than in the control group (**Fig. 2D-E**), suggesting that NOLC1 enhance the Cis-resistance in GC cells.

**Fig. 2.**
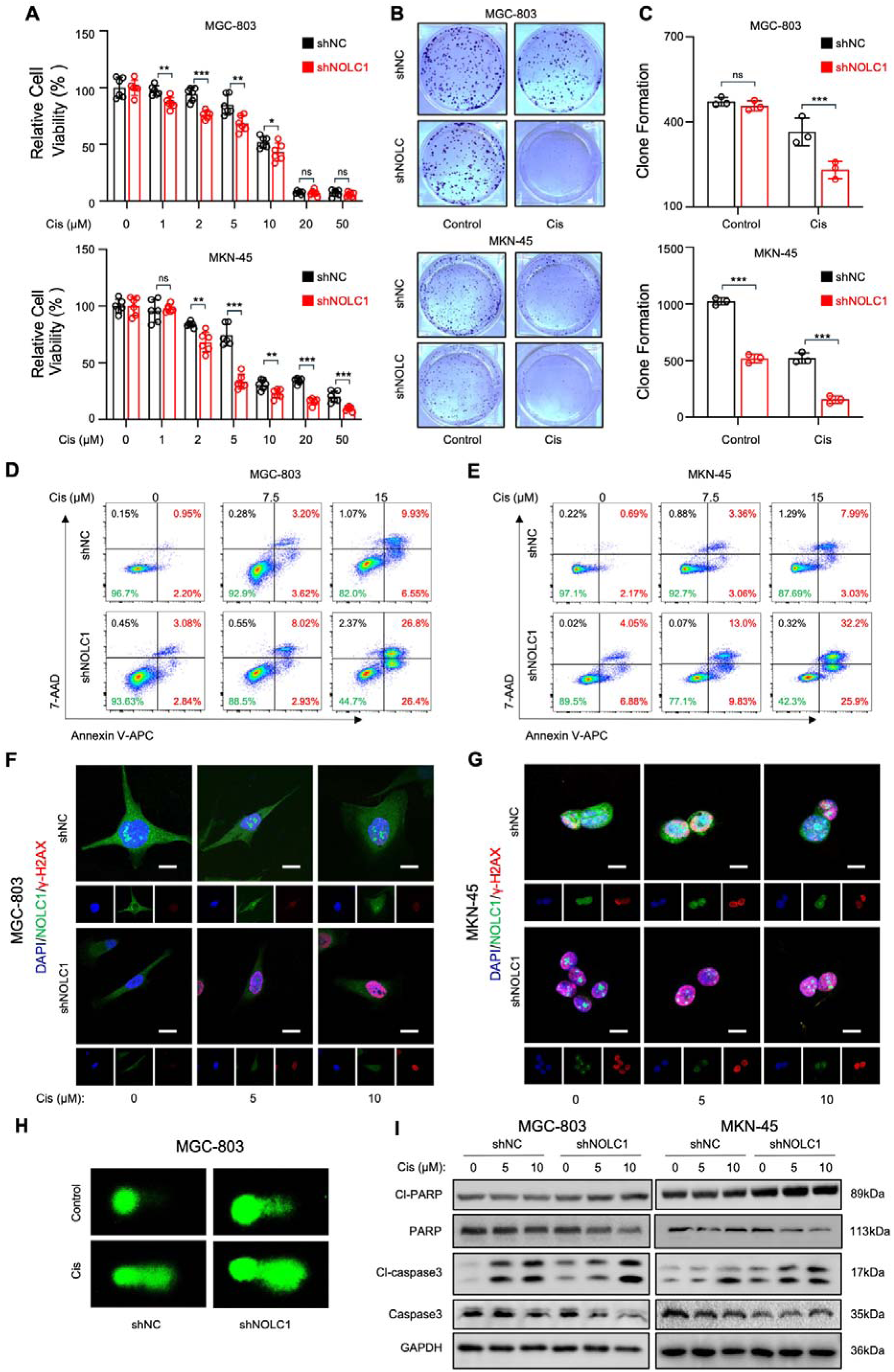
NOLC1 promoted Cis resistance in GC. **A** CCK-8 assay of GC cells transduced with shNC or shNOLC1 lentivirus and treated with different concentrations of Cis (*n* = 6). **B, C** Colony formation assay of GC cells transduced with shNC or shNOLC1 lentivirus and treated with PBS or Cis (15 μM). (B) Representative images of the colony formation assay. (C) Quantitative analysis of colony formations assay (*n* = 3). **D, E** Annexin V-APC and 7-AAD staining of GC cells transduced with shNC or shNOLC1 lentivirus and treated with the indicated concentrations of Cis, as analyzed via FACS. (D) MGC-803 cells and (E) MKN-45 cells. **F, G** Immunofluorescence staining assays of γ-H_2_AX in GC cells transduced with shNC or shNOLC1 lentivirus and treated with the indicated concentrations of Cis; (F) MGC-803 cells; scale bar = 10 μm. (G) MKN-45 cells; scale bar = 5 μm. **H** Comet assay of MGC-803 cells transduced with shNC or shNOLC1 lentivirus and treated with PBS or Cis (15 μM). **I** WB results of cleaved PARP, PARP, cleaved caspase-3, and caspase-3 in GC cells transduced with shNC or shNOLC1 lentivirus and treated with indicated concentrations of Cis. The data are presented as the means ± SDs. ns, nonsignificant; *p < 0.05; **p < 0.01; ***p < 0.001.

Since Cis exerts its cytotoxic effects by binding to DNA to form DNA-platinum adducts and destroy DNA^30^. The toxic effects of Cis on DNA were further evaluated via immunofluorescence (IF) staining of γ-H2AX (Ser-139) and comet assays. The IF results revealed increased aggregation of γ-H2AX foci in the knockdown group after treatment with Cis (**Fig. 2F-G**). Comet assays involving electrophoresis were performed and revealed that negligible DNA fragments migrated out of the nucleus in the negative control (NC) group treated with phosphate-buffered saline (PBS), while DNA tails were observed outside the nucleus after treatment with Cis (**Fig. 2H**). In particularly, a more pronounced tailing shape of DNA strands was found in the knockdown group, especially after treatment with Cis, indicating that NOLC1 knockdown enhances Cis-induced DNA damage. Moreover, the WB results revealed that the protein levels of cleaved caspase-3 and cleaved PARP were increased after treatment with Cis, especially in the knockdown group, which presented a greater degree of increasing accompanied by decreased expression of PARP and caspase-3 (**Fig. 2I, Supplementary Fig. S5B**) suggesting that Cis induced more severe DNA damage in the NOLC1 knockdown group. Taken together, these findings indicate that NOLC1 knockdown enhances Cis cytotoxicity in GC cells.

To determine whether NOLC1 affects GC Cis-resistance *in vivo*, we constructed xenograft GC models in BALB/c-nu mice. Randomly divided the mice into two groups and subcutaneously injected with MGC-803 cells (shNC or shNOLC1). When the tumor volume reached about 100 mm^3^, randomly divided each group into two subgroups (PBS and Cis), for a total of four groups. Cis or PBS was then administered to the mice via intraperitoneal injection twice per week for a total of 6 times (**Fig. 3A**). The results showed that Cis treatment effectively slowed the growth of the xenografts. The NOLC1-knockdown groups were more sensitive to Cis (**Fig. 3B-E**). Next, we further compare the therapeutic effects used hematoxylin and eosin (H&E) and IHC staining to detect the expression of cleaved caspase-3 and Ki-67 in the tumors. H&E staining revealed that the NOLC1-knockdown groups presented a larger necrotic area in the tumor tissues after Cis treatment (**Fig. 3F-G**). The IHC results revealed that Cis decreased Ki-67 protein levels, and the NOLC1-knockdown groups presented lower levels than did the NC group. The cleaved caspase-3 level was increased after Cis treatment, and in the knockdown groups, the cleaved caspase-3 level was greater than that in the NC groups. Collectively, all these data indicate that NOLC1 knockdown increased Cis sensitivity in gastric xenografts *in vivo*.

**Fig. 3.**
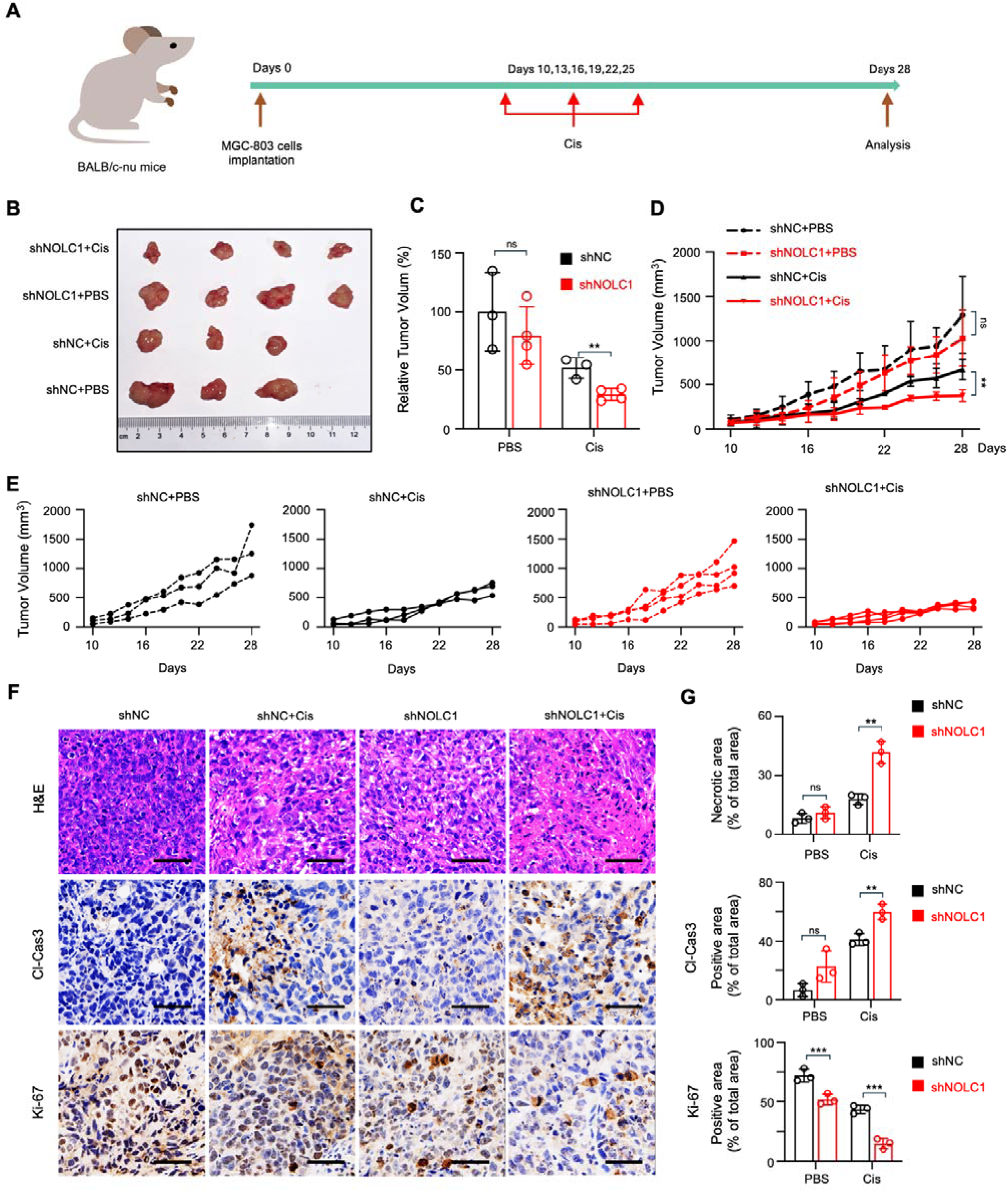
NOLC1 promoted Cis resistance *in vivo*. **A** Schematic representation showing the treatment of the subcutaneous xenograft tumor models. **B** Photographs of the corresponding tumors after indicated treatments. **C** Relative tumor volume after different treatments. **D, E** Tumor growth curves after indicated treatments. **F** H&E staining and IHC analysis of cleaved caspase-3 and Ki-67; scale bar = 50 μm. **G** Quantification analysis of the necrotic area and cleaved caspase-3 and Ki-67 expression in tumor tissue from mice given the indicated treatments (*n* = 3). The data are presented as the means ± SDs. ns, nonsignificant; *p < 0.05; **p < 0.01; ***p < 0.001.

### NOLC1 suppresses Cis-induced ferroptosis

Cis can induce apoptosis to kill cancers^31^. Also, other functions of Cis, such as inducing ferroptosis, which plays a pivotal role in Cis resistance have also been reported^32,33^. To determine which type of regulated cell death (RCD) is primarily blocked by NOLC1, we used transmission electron microscopy (TEM) to observe specific morphological changes in the MGC-803 and MKN-45 cells. The results revealed that cells with NOLC1 knockdown exhibied shrunken mitochondria and increased membrane density. After Cis treatment, the mitochondrial membrane was severely disrupted, and the mitochondrial cristae disappeared in the knockdown group (**Fig. 4A, Supplementary Fig. S6A**). These are the classic characteristic morphological features of ferroptosis. Moreover, the erastin (a ferroptosis activator) induced cell death was also suppressed by NOLC1 (**Supplementary Fig. S6B**). Fer-1 (a ferroptosis inhibitor) could rescue the sensitization effect of NOLC1 knockdown (**Supplementary Fig. S6C**). These findings indicate that NOLC1 might strongly inhibit ferroptosis to promote Cis resistance. During ferroptosis, ferroptotic cell lipid membrane structure is disrupted. We further examined specific ferroptotic cell lipid membrane morphological changes in response to Cis after NOLC1 knockdown. Lactate dehydrogenase (LDH) release assays revealed that after treatment with Cis, higher levels of LDH were detected in the knockdown groups (**Fig. 4B, Supplementary Fig. S6D**), indicating more severe destruction of the membrane structure in the knockdown group. Meanwhile, the high level of LDH release could be blocked by Fer-1 in knockdown groups, further reflected that NOLC1 promotes Cis resistance by suppresses ferroptosis (**Supplementary Fig. S6E**). Moreover, the JC-1 probe results indicated that the mitochondrial membrane potential decreased in the NOLC1 knockdown group after Cis treatment, indicating that the mitochondrial membrane was also disrupted (**Fig. 4C, Supplementary Fig. S6F-G**). These data revealed that NOLC1 knockdown GC cells displayed classic ferroptosis-related morphological features.

**Fig. 4.**
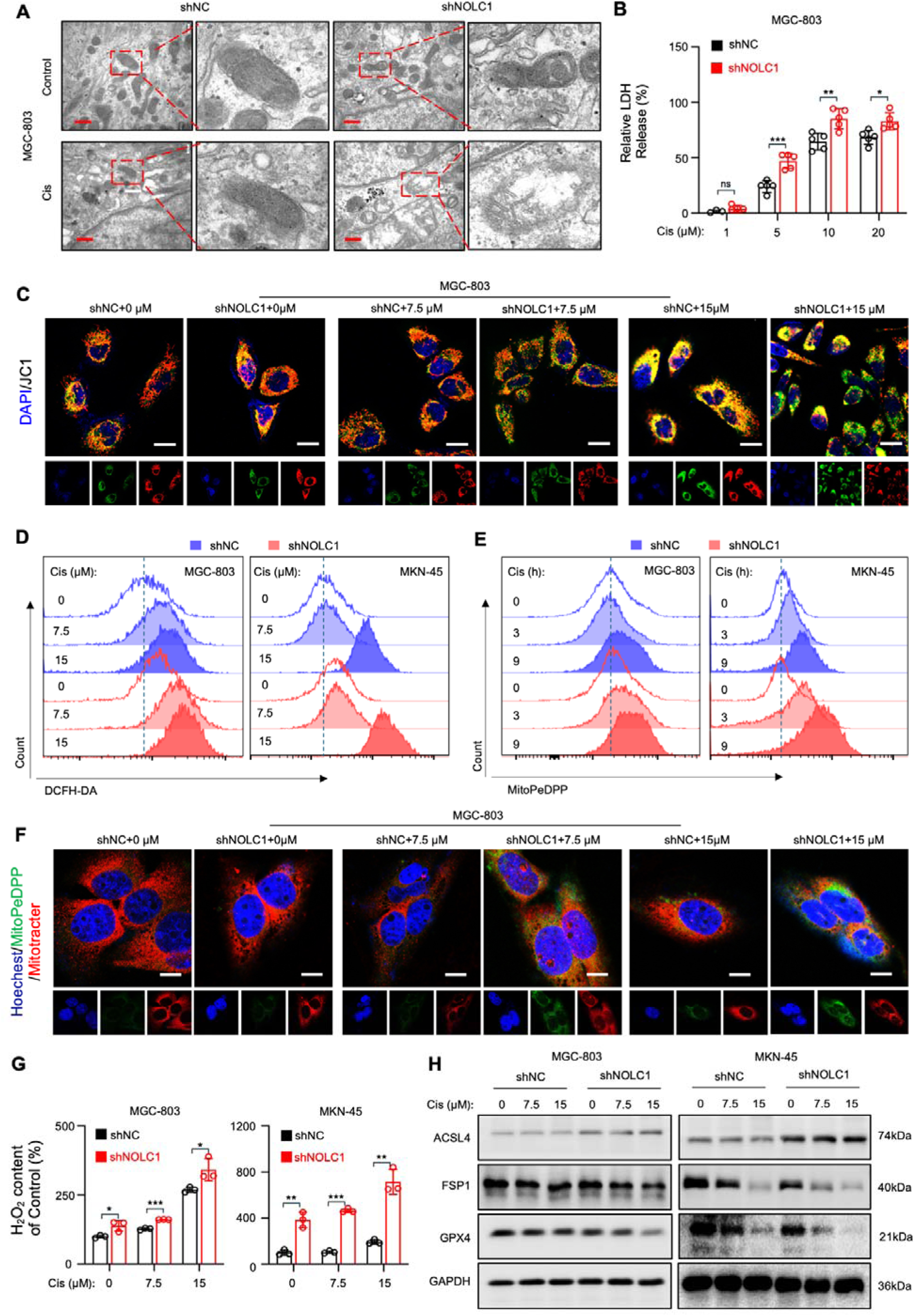
NOLC1 deletion rendered GC susceptible to ferroptosis. **A** TEM image of MGC-803 cells transduced with shNC or shNOLC1 lentivirus and treated with PBS or Cis (15 µM); scale bar = 400 nm. **B** LDH release analysis of MGC-803 cells transduced with shNC or shNOLC1 lentivirus and treated with indicated concentrations of Cis (*n* = 5). **C** Representative JC-1 fluorescence images of MGC-803 cells transduced with shNC or shNOLC1 lentivirus and treated with indicated concentrations of Cis; scale bar = 20 µm. **D** Fluorescence intensity of DCFHA-DA measured by FACS in GC cells transduced with shNC or shNOLC1 lentivirus and treated with indicated concentrations of Cis. **E** Fluorescence intensity of MitoPeDPP measured by FACS in GC cells transduced with shNC or shNOLC1 lentivirus and treated with Cis (15 µM) for different time. **F** Representative MitoPeDPP fluorescence images of MGC-803 cells transduced with shNC or shNOLC1 lentivirus and treated with indicated concentrations of Cis; scale bar = 10 µm. **G** Relative H_2_O_2_ content of GC cells transduced with shNC or shNOLC1 lentivirus and treated with indicated concentrations of Cis (*n* = 3). **H** WB result of ACSL4, FSP1 and GPX4 protein levels in GC cells transduced with shNC lentivirus or shNOLC1 lentivirus and treated with different concentrations of Cis. The data are presented as the means ± SDs. ns, nonsignificant; *p < 0.05; **p < 0.01; ***p < 0.001.

Disruption of the cell lipid membrane during ferroptosis is caused by lipid peroxides^8^. Therefore, we further detected the reactive oxygen species (ROS) level after NOLC1 knockdown using 2,7-dichlorodihydrofluorescein diacetate (DCFH-DA) staining and analyzed by fluorescence activated cell sorting (FACS) and confocal microscopy. Compared with the negligible ROS accumulation of the NC groups, the knockdown groups presented a greater level of ROS accumulation, especially when treated with Cis (**Fig. 4D, Supplementary Fig. S6H-I**). MitoPeDPP, which accumulates in the inner mitochondrial membrane, can be oxidized by lipid peroxides and release strong fluorescence. Using MitoPeDPP probes, we found that, in the NOLC1-knockdown groups, the rate of mitochondrial-specific ROS accumulation was significantly higher than that in the negative control (NC) group at the same Cis treatment time. (**Fig. 4E, Supplementary Fig. S6J**). Also, a greater level of fluorescence accumulation in the mitochondria was observed after treatment with Cis in the knockdown groups (**Fig. 4F**). Moreover, the H_2_O_2_ content increased after Cis treatment, and the H_2_O_2_ content in the knockdown groups was much greater than that in the NC groups (**Fig. 4G**). These data indicated that lipid ROS significantly accumulated in the NOLC1 knockdown group. In line with the above data, the key protein involved in ferroptosis, GPX4, were obviously downregulated in the NOLC1 knockdown group both *in vitro* and *in vivo* (**Fig. 4H, Supplementary Fig. S7A, B**). Moreover, FSP1 was decreased and ACSL4 was increased in the knockdown group reflecting that NOLC1 could effectively suppress ferroptosis. Taken together, these data show that NOLC1 primarily inhibits Cis induced ferroptosis to promote GC resistance.

### NOLC1 binds to p53 and blocks p53 translocation

Next, we further investigated the molecular mechanism of which NOLC1 promotes resistance in GC. Recent research has shown that the function of NOLC1 is associated with p53^28,29^. In addition, via mRNA-seq, we found that the p53 signal pathway was obviously altered in MGC-803-CR cells (**Supplementary Fig. S3E**). Thus, we assumed that NOLC1-mediated suppression of ferroptosis promotes GC resistance via regulating p53.

To verify our hypothesis, the p53 protein level was measured, and the results indicated that the p53 protein level increased when the cells were treated with Cis, further confirming that p53 is upregulated in stressed cells (**Fig. 5A, Supplementary Fig. S8A**). However, in the knockdown group, the p53 level was lower than that in the NC group and was not associated with cell death trends. BAX is a proapoptotic protein whose transcription is mediated by p53^34^. BCL-2 is an antiapoptotic protein which is mediated by p53 via protein interaction in cytoplasm. Intersetingly, the increase of BAX level after NOLC1 knockdown was out of line with the p53 protein level trend but in line with the cell death trend, but the BCL-2 level was in line with p53 level (**Fig. 5A, Supplementary Fig. S8B**). Additionally, according to the IHC results, p53 expression was also decreased in the knockdown group *in vivo* (**Supplementary Fig. S8C**). Thus, we assume that after NOLC1 is knocked down, the transcriptional functions of p53 increased, but the protein level decreased. To validate our hypothesis, we detected the transcriptional activity levels of p53 and downstream genes of p53 (CDKN1A, BAX, FAS, and PTEN) expression. Dual luciferase assay shown p53 transcription is activated by Cis, indicating that p53 is activated when cells are stressed. Knocking down NOLC1 significantly increased p53 activity in both MGC-803 and MKN-45 cells (**Fig. 5B, Supplementary Fig. S8D)**. Meanwhile, NOLC1 overexpression in HEK-293T cells obviously decreased p53 transcriptional activity in a dose-dependent manner (**Fig. 5C**). Additionally, the activity of NC reporter and that of the mut p53 reporter did not significantly differ (**Supplementary Fig. S8E, F**). Real-Time quantitative reverse transcription polymerase chain reaction (RT-qPCR) results revealed that CDKN1A, BAX, FAS, and PTEN mRNA levels were increased in the knockdown group both *in vitro* and *in vivo* (**Supplementary Fig. S8G, H**). Collectively, these results indicate that NOLC1 can decrease the transcriptional activity of p53.

**Fig. 5.**
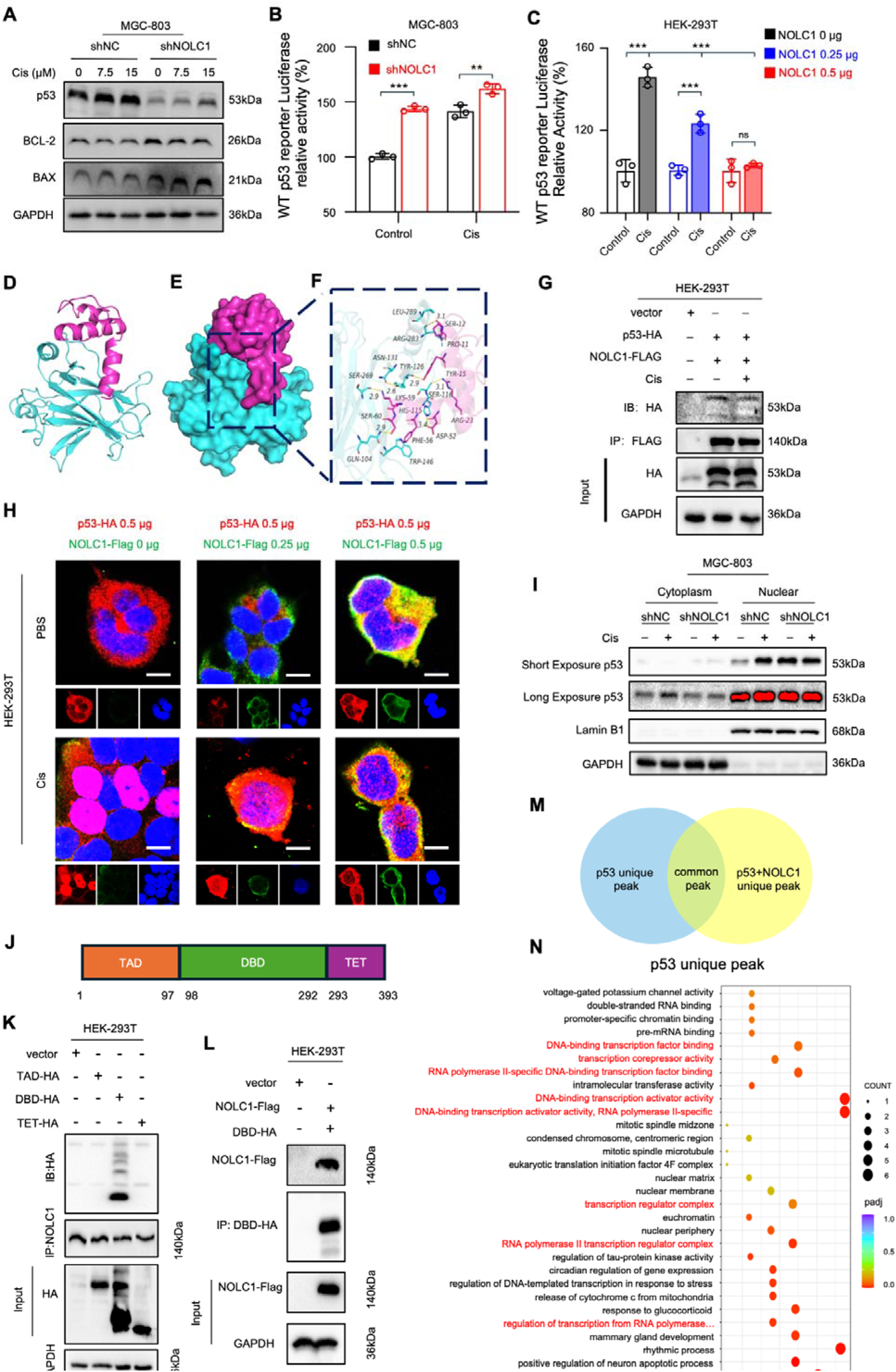
NOLC1 decreased p53 transcriptional activity. **A** WB analysis of p53 BCL-2 and BAX in MGC-803 cells transduced with shNC or shNOLC1 lentivirus and treated with indicated concentrations of Cis. **B**, **C** Luciferase assay of p53 transcription activity (B) in MGC-803 cells transduced with shNC or shNOLC1 lentivirus and treated with PBS or Cis (15 µM) and (C) in HEK-293T cells transfected with 0.00, 0.25, or 0.50 µg NOLC1 plasmid and treated with PBS or Cis (15 µM). **D**D**F** Docking analysis of NOLC1 and p53. (D) The backbones of the proteins were shown in tubes and colored red (NOLC1) and cyan (p53). (E) The NOLC1 and p53 proteins were shown on the surface. (F) The detailed binding mode of NOLC1 with p53. The yellow dash represented the hydrogen bond. **G** Co-IP assay of NOLC1 with p53. **H** Representative NOLC1-Flag and p53-HA fluorescence images of HEK-293T transfected with indicated amount of NOLC1-Flag and p53-HA plasmids, and treated with PBS or Cis (15 µM); scale bar = 10 µm.. **I** WB analysis of p53 protein levels in the cytoplasm or nucleus of MGC-803 cells transduced with shNC or shNOLC1 lentivirus and treated with PBS or Cis (15 µM). **J** Schematic map of the p53 functional domain. **K** Co-IP assay of NOLC1 with different p53 functional domains. L Co-IP assay of p53 DBD-HA with NOLC1-Flag. **M** Schematic map of the ChIP-seq analysis. **N** GEO analysis of the unique p53 peak identified via ChIP-seq. The data are presented as the means ± SDs. ns, nonsignificant; **p < 0.01; ***p < 0.001.

Considering that, as a molecular chaperone, NOLC1 can interact with other proteins, such as TRF2^35^, to regulate their functions, we assume that NOLC1 regulates p53 functions via interacts with p53. Thus, the docking studies were performed. The results showed that NOLC1 can interact with p53 (**Fig. 5D**). The binding energy of NOLC1 to p53 was -236.13 kcal/mol, and the binding sites of p53 are all in the DNA-binding domain (DBD), which is responsible for transcription (**Table 1**). In addition, the surface of the p53 protein matched well with that of the NOLC1 protein, which promoted stable binding (**Fig. 5E, F**). To verify the potential interaction between NOLC1 and p53, we carried out coimmunoprecipitation (Co-IP) assays in HEK-293T cells. As expected, immunoprecipitation of p53-HA coprecipitated with NOLC1-Flag demonstrated that NOLC1 can interact with p53 (**Fig. 5G**). The IF results also revealed that NOLC1 colocalized with p53 in both HEK-293T and MGC-803 cells (**Supplementary Fig. S9A**). The nuclear accumulation of p53 is crucial for its transcriptional function. To detected whether NOLC1 effect the p53 nuclear accumulation upon Cis treatment, IF was applied to detect the p53 location after Cis treatment in HEK-293T cells. Under normal conditions, NOLC1 and p53 colocalized in the cytoplasm, and p53 accumulated into the nuclear after Cis treatment. After overexpression of NOLC1, the nuclear level of p53 significantly decreased under Cis treatment, and as NOLC1 expression increased, the nuclear/cytoplasmic ratio of p53 gradually decreased further. (Fig 5H, Supplementary **Fig. S9B**). In addition, knockdown of NOLC1 significantly increased the nuclear/cytoplasmic ratio of p53 under normal conditions **(Supplementray Fig. S9C**). The WB results revealed similar trends in the GC cells (**Fig. 5I**). Additionally, IHC staining revealed that the ratio of p53 nuclear positive cells was significantly increased (**Supplementary Fig. S8C**). These data indicate that NOLC1 inhibits p53 nuclear accumulation.

**Table 1.** Docking results of the two target proteins.

| Protein1 | Protein2 | Binding Energy<br>(kcal/mol) | Contact Sites<br>(protein1) | Contact Sites<br>(protein2) | Combination<br>Type |
| --- | --- | --- | --- | --- | --- |
| NOLC1 | TP53 | -236.13 | SER-60, SER-12,<br>LYS-59, TYR-15,<br>ASP-52, ARG-23,<br>PRO-11, PHE-56 | GLN-104, ARG-283,<br>ASN-131, SER-269,<br>TYR-126, HIS-115,<br>SER-116, LEU-289,<br>TRP-146 | Hydrogen bond,<br>Hydrophobic interaction |

The full-length p53 protein has three conserved functional domains: a transactivation domain (TAD), a DNA binding domain (DBD), and a tetramerization domain (TAE)^36^ (**Fig. 5J**). To determine which domain of p53 is responsible for its interaction with NOLC1, three functional domain constructs were generated to perform Co-IP assays. Consistent with the docking results, Co-IP assays revealed that only the DBD can interact with NOLC1 (**Fig. 5K**). Meanwhile, the DBD-HA could coprecipitated NOLC1-Flag too (**Fig. 5L**). Considering that the interaction between NOLC1 and p53 might block p53 binding to DNA, we performed ChIP-Seq to determine whether this interaction could inhibit p53 binding to specific DNA sequences. We transfected HEK-293T cells with a p53 full-length plasmid or the p53 full-length and NOLC1 plasmid (**Fig. 5M**). Then, p53 immunoprecipitation confirmed the binding of p53 to the DNA sequence. As shown in **Fig. 5N**, the p53 group could uniquely bind to DNA-binding transcription activator activity peaks, DNA-binding transcription factor binding peaks, etc. These findings demonstrated that NOLC1 could inhibit p53 binding to transcription-related DNA sequence. Furthermore, the data indicated that NOLC1 binds to the p53 DBD, not only to mediate p53 nuclear translocation but also to block p53 binding to DNA, eventually blocking p53-mediated ferroptosis.

Taken together, the above data showed that NOLC1 inhibits p53 nuclear translocation stimulated by Cis treatment and transcriptional activity by interacting with the p53 DBD.

### NOLC1 inhibits p53 degradation via disruption of the MDM2**D**p53 feedback loop

Although we have shown that NOLC1 decreases p53 transcriptional functions, it is still unclear why the p53 protein level decreases after NOLC1 are knocked down (**Fig. 5A**). RT□qPCR results shown there was no significant difference of the p53 mRNA level between the NC and knockdown groups (**Fig. 6A**). Thus, we speculated that NOLC1 increases the p53 protein level by inhibiting its degradation. The E3 ubiquitin ligase MDM2, encoded by a p53-responsive gene, is the master antagonist of p53 by promoting p53 ubiquitination and proteasomal degradation, forming a feedback loop^37^. On the basis of the above data, we hypothesized that NOLC1 inhibits p53 transcription-mediated MDM2, thereby reducing p53 ubiquitination and degradation. Therefore, we next detected the MDM2 mRNA and protein levels. As is shown in **Fig. 6B-D**, the MDM2 mRNA and protein levels were both increased after Cis treatment, and in the knockdown groups, the levels were greater than those in the NC groups, indicating that NOLC1 knockdown also promoted p53 transcription to MDM2. Similarly, the protein level of MDM2 *in vivo* was also elevated in the knockdown group, as detected by IHC (**Fig. 6E, F**). We next examined the p53 turnover rate. In the OE-NOLC1 group, the p53 turnover rate was much slower than that in the vector group (**Fig. 6G**). Furthermore, ubiquitination assays indicated that NOLC1 inhibits p53 ubiquitination and that Cis treatment induce a marked decrease in the p53 ubiquitination level both in MGC-803 and HEK-293T cells (**Fig. 6H**). Collectively, these data indicate that NOLC1 inhibits p53 degradation by blocking the nuclear entry and transcriptional activity of p53. Therefore, the p53-MDM2 negative feedback loop is interrupted.

**Fig 6.**
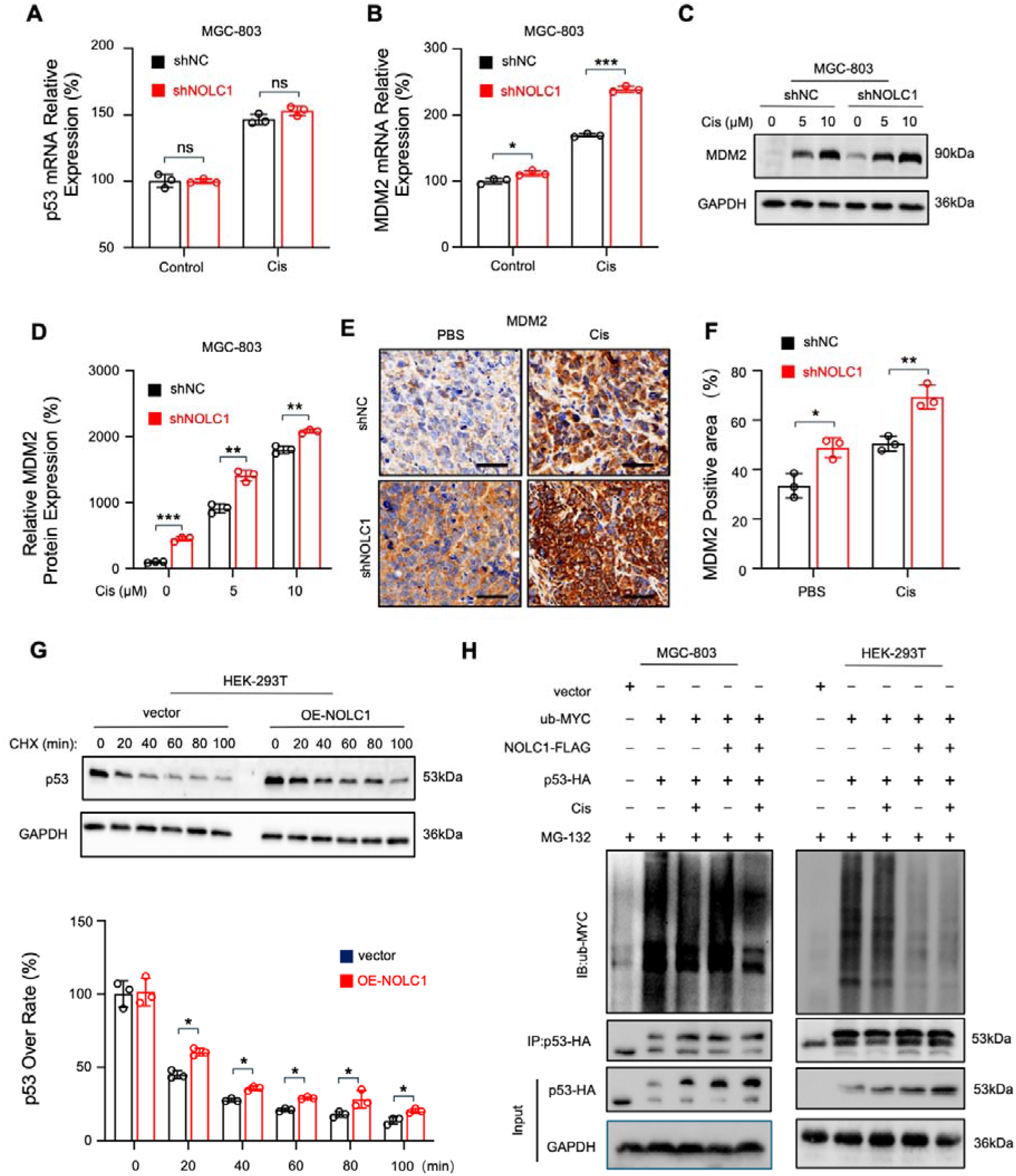
NOLC1 inhibited p53 ubiquitination and degradation. **A** RT□qPCR analysis of p53 mRNA in MGC-803 cells transduced with shNC or shNOLC1 lentivirus and treated with PBS or Cis (15 µM). **B** RT□qPCR analysis of MDM2 mRNA in MGC-803 cells transduced with shNC or shNOLC1 lentivirus and treated with PBS or Cis (15 µM). **C, D** WB assay of MDM2 in MGC-803 cells transduced with shNC or shNOLC1 lentivirus and treated with PBS or Cis (15 µM). (C) Representative images and (D) relative MDM2 protein levels. **E, F** IHC staining of MDM2 in MGC-803 tumors from BALB/c-nu mice. (E) Representative images; scale bar = 50 µm. (F) MDM2-positive area in MGC-803 tumors. **G** p53 turnover rate analysis in HEK-293T cells transfected with vector or NOLC1 plasmids and treated with CHX (40 µM) for the indicated times. **H** Ubiquitination assay in MGC-803 and HEK-293T cells transfected with the indicated plasmids. The data are presented as the means ± SDs. ns, nonsignificant; *p < 0.05; **p < 0.01;

### p53 knockdown rescued the sensitizing effect of NOLC1 knockdown

Given that NOLC1 can promote GC resistance and interact with p53 to inhibit p53 transcription, whether NOLC1 promotes resistance by inhibiting p53 functions is still unclear. Thus, we next used siRNA to knock down the p53 protein in MGC-803 cells. An Annexin V-APC/7-AAD apoptosis assay revealed that NOLC1 single-knockdown cells were more sensitive to Cis than were NC cells. However, the percentage of death cells in the p53/NOLC1 double-knockdown group was not significantly different from that in the NC group **(Supplementary Fig. S10A-B**). Furthermore, CCK-8 assay showed similar results (**Supplementary Fig. S10C**). These data show that NOLC1 promotes Cis resistance via mediating p53 functions. Moreover, compared to the NOLC1 single-knockdown group, the GPX4 protein level was increased in the p53/NOLC1 double-knockdown group (**Supplementary Fig. S10D**). These data demonstrates that NOLC1 inhibits ferroptosis via regulating p53.

### NOLC1 knockdown increases the efficacy of anti-PD-1 combined with cisplatin

Considering ferroptosis can promote ICD by releasing of DAMPs^14^. Herein, we further investigated whether NOLC1 knockdown can increase ferroptosis-mediated ICD. First, DAMPs, including LDH, high mobility group box-1 (HMGB1), and calreticulin (CRT), were analyzed further. As mentioned above, the release of LDH significantly increased in the NOLC1 knockdown group after Cis treatment (**Fig. 4B, Supplementary Fig. S6D**). Consistently, the fluorescence intensity of HMGB1 in the knockdown group was markedly lower than that in the NC group, especially when cells were treated with Cis (**Supplementary Fig. S11A, B**). These results confirmed that a large amount of HMGB1 was released into the extracellular environment after Cis treatment in the knockdown groups. Furthermore, the CRT fluorescence intensity in the knockdown groups were greater than that in the NC groups, especially treated with Cis (**Supplementary Fig. S11C, D**). These data showed that NOLC1 repressed ferroptosis-mediated ICD.

To further assess the role of NOLC1 in ferroptosis-mediated immunotherapy, subcutaneous mouse forestomach carcinoma (MFC) tumor-bearing 615 mice model were established for further study. Four-week-old male 615 mice were randomly divided into two groups and were subcutaneously injected with MFC cells (shNC or shNOLC1). When tumor volume reached about 100 mm^3^, each group was randomly divided into four subgroups (PBS, anti-PD-1, Cis, and Cis combined with anti-PD-1), for a total of eight groups. On the 7th, 10th, and 13th days, the mice were intraperitoneal injected drugs (Cis, anti-PD-1 antibody) **(Fig. 7A)**. As shown in **Fig. 7B-C, and Supplementary Fig. S12A**, anti-PD-1 monotherapy had no significant effect in the presence of the immunosuppressive TME. Cis treatment effectively blocked the growth of tumors. PD-1 combined with Cis had the best efficacy. Importantly, NOLC1 silencing obviously increased the efficacy of Cis alone and anti-PD-1 plus Cis. Accordingly, the tumor weight data showed similar trends (**Fig. 7D**). Subsequently, the therapeutic effect was confirmed by H&E and IHC staining. H&E staining revealed the same trend: the NOLC1 knockdown group treated with Cis+anti-PD-1 presented the most severe necrosis in the tumor tissues (**Supplementary Fig. S12B, C**). As shown in **Fig. 7E, and Supplementary S12D-F**, Cis therapy increased the cleaved caspase-3 level and decreased the Ki-67and GPX4 levels. The Cis plus PD-1 group presented more significant changes than the Cis group did. Compared with those in the NC groups, the NOLC1 knockdown groups had higher cleaved caspase-3 and lower GPX4 and Ki-67 expressions. These findings demonstrated that NOLC1 knockdown increased the efficacy of the Cis monotherapy and combination treatment. Additionally, the knockdown group treated with anti-PD-1 plus Cis presented the highest level of nuclear p53 staining (**Supplementary Fig. S12G**), indicating that NOLC1 also mediates p53 nuclear accumulation in the MFC tumor model. Liperfluo is specifically oxidized by lipid peroxides and emits strong fluorescence. Liperflou staining revealed that the NOLC1-knockdown groups treated with Cis plus PD-1 presented the most intense fluorescence (**Fig. 7E**). The HMGB1 level in the serum was also increased in the knockdown group treated with anti-PD-1 plus Cis (**Supplementary Fig. S12H**). Indicating NOLC1 suppresses ferroptosis in MFC tumors. Collectively, these data suggest that NOLC1 decreases the antitumor efficacy of anti-PD-1 plus Cis therapy via suppressing Cis-induced ferroptosis.

**Fig 7.**
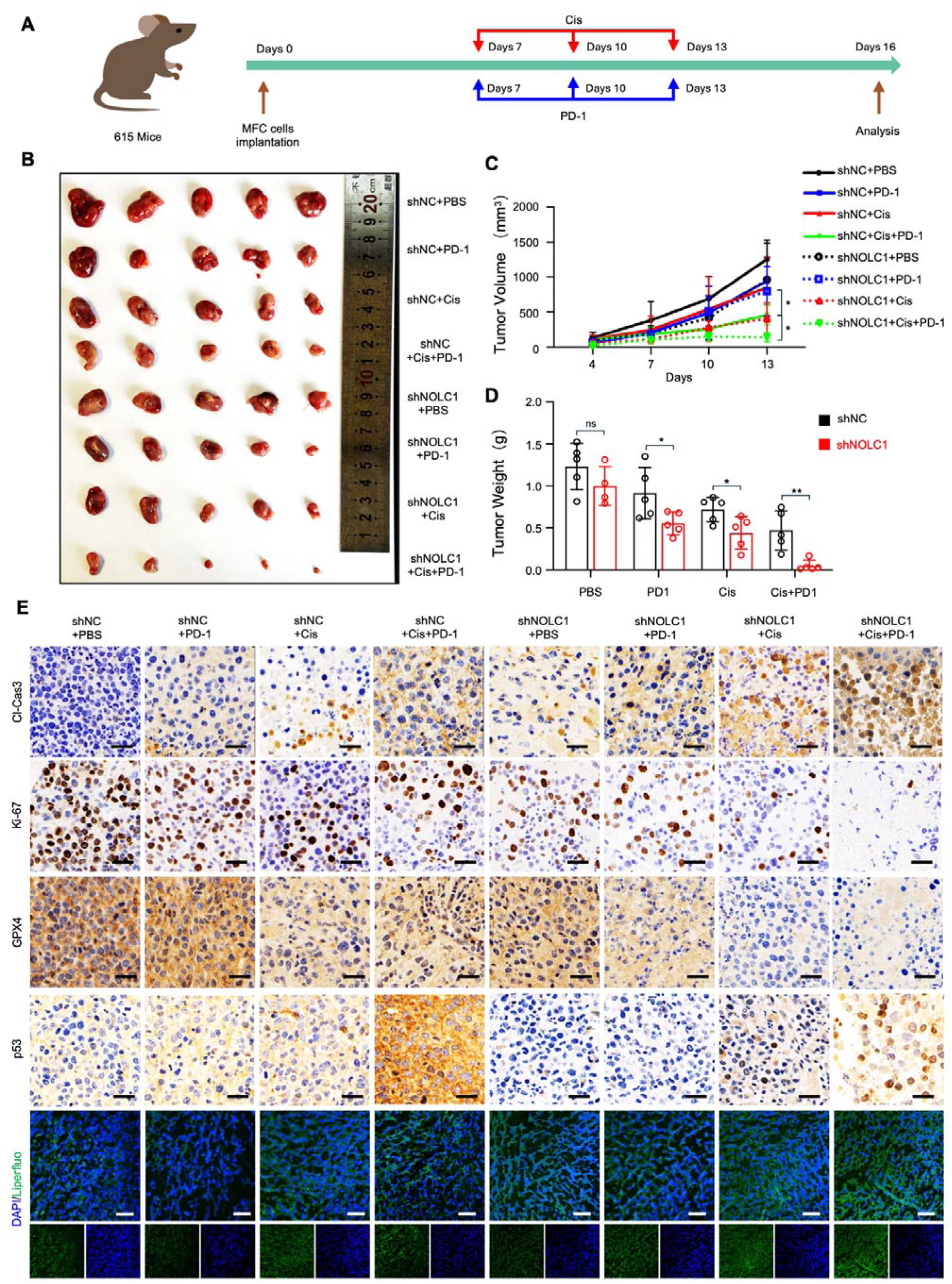
NOLC1 knockdown increased the efficacy of Cis combined with anti-PD-1 treatment. **A** Schematic representation showing the treatment of the subcutaneous tumors. **B** Corresponding tumor photographs after different treatments. **C** Tumor growth curves after different treatments. **D** Tumor weight after the indicated treatment. **E** Representative images of Liperfluo staining and IHC analysis of cleaved caspase-3, Ki-67, GPX4, and p53 in tumor tissues; IHC analysis scale bar = 25 μm, liperfluo staining scale bar = 50 μm. The data are presented as the means ± SDs. ns, nonsignificant; *p < 0.05; **p < 0.01.

### Silencing NOLC1 promotes reprogramming of the TME by Cis combined with anti-PD-1 immunotherapy

The tumor microenvironment (TME) plays an important role in GC progression and therapeutic outcome and is correlated with the patient response to immunotherapy^38^. In addition, ferroptosis can also reprogram the TME to increase immunotherapy efficacy^39^. Therefore, we next explored the effect of NOLC1 knockdown on the TME. First, we collected blood samples and extracted lymphocytes. FACS was subsequently used to measure the abundance of specific lymphocytes. As is shown in **Fig. 8A** the proportions of CD3^+^CD8^+^ cytotoxic T lymphocytes (CTLs), which have been reported to play pivotal roles in killing cancer cells^40^, are significantly increased after treated with Cis plus anti-PD-1, and the proportions of CTLs in the knockdown group are much higher than that in the NC group. In addition, Cis monotherapy could increase the proportion of CTLs in the knockdown group, but not in the NC group. Furthermore, the IF staining results also revealed an increase in the number of CTLs infiltration in tumors in the knockdown group, especially in those treated with Cis plus anti-PD-1(**Fig. 8B**). The levels of tumor necrosis factor-_a_ (TNF-_a_), interferon-_y_ (IFN-_y_), and interleukin 6 (IL-6) in the combination treatment groups were significantly increased, and monotherapy did not effectively increase the levels of these factors (**Fig. 8C**). In the knockdown groups, this activation was much more obvious. Taken together, these data demonstrate that NOLC1 knockdown can promote ferroptosis-mediated ICD and reprogram the TME.

**Fig 8.**
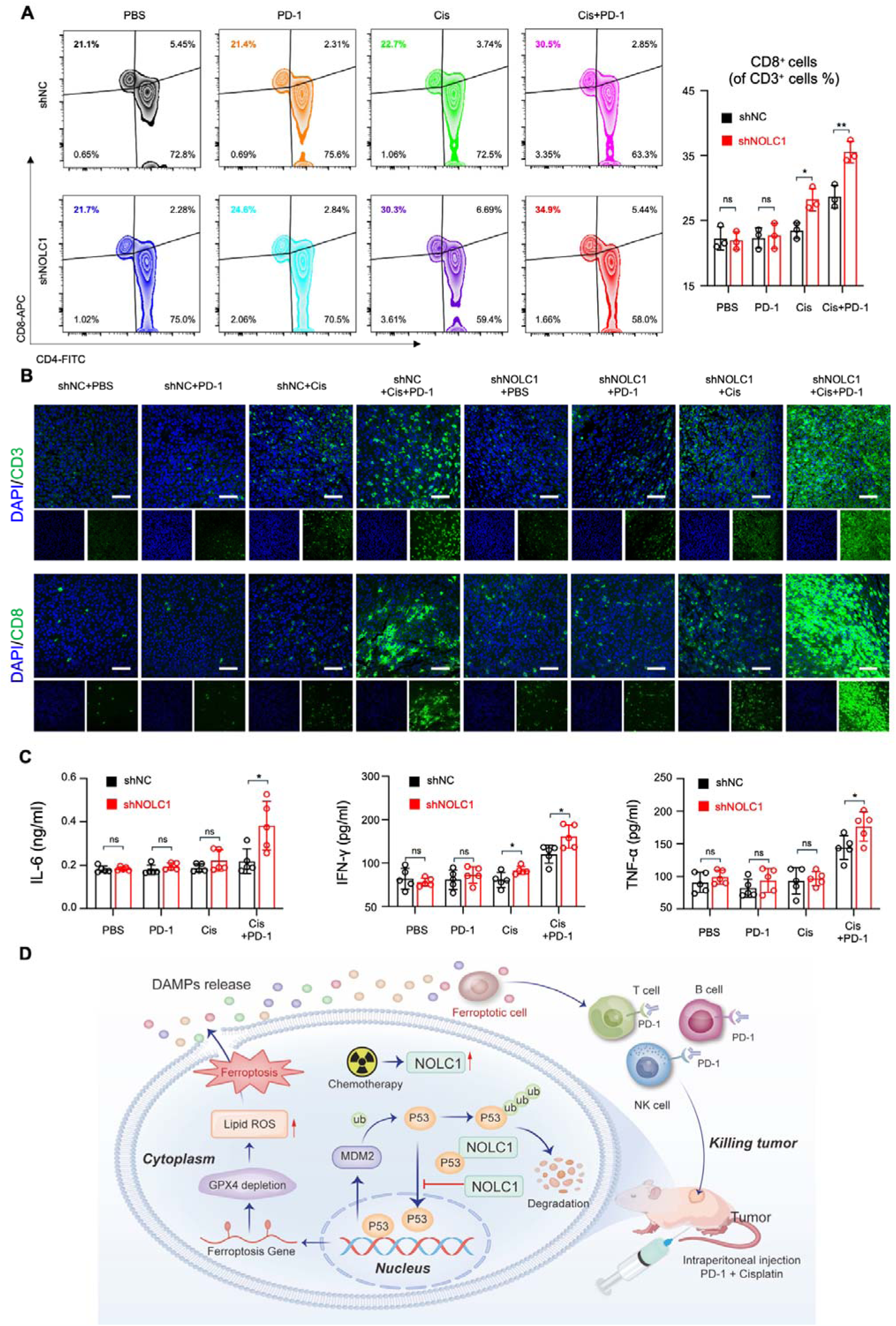
Knockdown of NOLC1 reprogrammed the tumor microenvironment after Cis and anti-PD-1 combination therapy. **A** FACS analysis of peripheral blood lymphocytes. **B** Representative immunofluorescence staining of CD-3 and CD-8 in MFC tumors after indicated treatments; scale bar = 50 µm. **C** The serum levels of inflammatory factors (IL-6, IFN-_y_, and TNF-_a_) (*n* = 5). **D** Schematic illustration of the mechanism of NOLC1 inhibits p53 mediate ferroptosis to increase the efficacy of anti-PD-1 plus Cis in GC. The data are presented as the means ± SDs. ns, nonsignificant; *p < 0.05; **p < 0.01.

### Biosafety of combined therapy with cisplatin and PD-1

Despite the efficacy of immunotherapy combined with chemotherapy, there are still many side effects due to the toxicity of immune checkpoint inhibitors or chemotherapeutic agents^41^.To further assess the biosafety of combination therap*y in viv*o, we collected major organs and blood from the mice for further analysis. Analysis of liver function markers, such as alanine transaminase (ALT) and aspartate aminotransferase (AST), and kidney function markers, such as blood urea nitrogen (BUN), indicated that no significant renal or hepatic toxicity was induced by the treatment (**Supplementary Fig. S14A**). The organs were assessed via H&E staining. Compared with PBS treatment, combination therapy did not induce significant morphological changes or inflammation in the heart, liver, spleen, lung, or kidney (**Supplementary Fig. S14B**). Collectively, these data indicate the satisfactory biocompatibility of the combination therapy.

## Discussion

Despite advances in GC therapy, chemotherapy resistance remains the major obstacle to improving patient outcomes^3^. In this study, we identified NOLC1 as a mediator of anti-PD-1 plus Cis resistance in GC. The functions of NOLC1 in cancer is controversial, as evidenced by its pro- or antitumor potential in different tumor types and models^27^. NOLC1 functions as an oncogene in various tumors via different pathways. In esophageal cancer (ESCA), NOLC1 can activate the PI3K-AKT signaling pathway^42^. In ovarian cancer (OC), circ-NOLC1 binds to ESRP1 and upregulates CDK1 and RhoA expression to promote OC progression^43^. More importantly, NOLC1 promotes non-small cell lung cancer resistance to multiple drugs. However, NOLC1 act as a tumor suppressor in clear cell renal cell carcinoma (ccRCC)^44^. The results of this study indicate that NOLC1 is up-regulated in GC tumors and Cis-resistant GC cells. In addition, silencing NOLC1 affected the efficacy of Cis and anti-PD-1 plus Cis combination treatment.

To determine its mechanism of action, we performed TEM imaging. The results revealed that after NOLC1 knockdown, cells displayed classical morphological features of ferroptosis, and treatment with a ferroptosis inhibitor counteract the Cis-sensitization effect of NOLC1 knockdown. These findings demonstrated that NOLC1 inhibits Cis-induced ferroptosis. Next, via mRNA-seq, we found that the p53 pathway, which is associated with NOLC1 functions, was significantly altered in MGC-803-CR cells. As a molecular chaperone, NOLC1 can interact with other proteins and regulate their translocation. We also revealed that NOLC1 interacts with p53 and inhibits p53 nuclear accumulation stimulated by Cis treatment rather than regulating p53 total protein levels, thus inhibiting p53 transcriptional activity. Eventually, p53-mediated ferroptosis is suppressed.

The mechanisms by which p53 promotes ferroptosis remain to be fully explored. Ferroptosis, a novel type of RCD, has received considerable attention in both basic and clinical research ^45^. Recently, studies have revealed that p53 can significantly promote ferroptosis via different mechanisms. For example, p53 could transcriptional and untranscriptional mediate SLC7A11, a key component of the cystine/glutamate antiporter^46,47^. Additionally, p53 can transcriptionally mediate several key ferroptosis proteins, such as TIGAR and GLS2^16^. In this study, we found that inhibiting p53 transcriptional functions could markedly suppress ferroptosis in GC cells. These findings indicate that the transcriptional functions of p53 are critical for mediating ferroptosis.

Protein□DNA interactions are central to the response of the tumor suppressor p53 to numerous stress signals. Under various stresses, p53 can enter the nuclear and bind to specific DNA via interactions between response elements (REs) and the p53 DBD^36^. Our results showed that NOLC1 interacts with the p53 DBD, which could block p53 binding to the DNA sequence, eventually inhibiting its tumor-suppressive functions. By using ChIP-seq, we found that NOLC1 strongly inhibited p53 binding to the transcription-associated DNA sequence (**Fig. 5M**). Our results show that the interaction between NOLC1 and p53 not only inhibits p53 binding to the DNA sequence but also decreases p53 nuclear accumulation. As shown in **Supplementary Fig. S8A**-**B**, the interaction between NOLC1 and p53 was limited to the cytoplasm rather than the nucleus. NOLC1-mediated inhibition of p53 functions relies mainly on blocking p53 nuclear accumulation rather than inhibiting its interaction with DNA.

Traditionally, the ability of tumor to inhibit p53 functions relies primarily on the MDM2□p53 negative feedback loop or mutant p53 functions^48^. Our results reveal a novel way to inhibit p53 functions: sequestration of the p53 protein in the cytoplasm. p53 is mutated in 50% of cancers, and most of these mutation occur in the DBD (86% of tumorigenic mutants are present in the DBD)^49,50^. Therefore, this suppressive effect of NOLC1 on p53 is limited to only WT p53, and this combination of p53 and NOLC1 could further inhibit p53 functions.

More and more studies have shed light on the important role of p53 in immunotherapy. The activation of p53 suppresses tumor immune evasion and promotes the antitumor immune response via the cGAS-STING pathway, regulating T-cell-mediated antitumor immunity, etc^51^. In addition, mut p53 exerts immunosuppressive effects via multiple mechanisms, such as increasing tumor-associated neutrophil infiltration, decreasing CTLs infiltration, and promoting fibrosis^52^. Our results demonstrated that activating p53 transcriptional functions could enhance the immune response and reprogram the TME by promoting ferroptosis. Current treatments for GC have evolved significantly through systemic therapies and include chemotherapy, targeted therapy and immunotherapy. Immunotherapy has substantially advanced the treatment of GC with microsatellite instability (MSI) but has limited efficacy for GC with microsatellite stability (MSS)^4^. However, MSI status is observed for only approximately 20% of GC patients^21,53^. Thus, identifying new molecular biomarkers is highly important. In this study, we found that NOLC1 knockdown could transform the chemoresistance tumor environment to a sensitive state. Thus, lower expression of NOLC1 could be a marker for identifying patients with a better response to combination treatment of anti-PD-1 plus Cis. Additionally, our studies identified NOLC1 could be promising target to increasing ICI sensitivity in MSS GC.

In conclusion, our study demonstrated that NOLC1 binds to the p53 DBD and inhibits p53 nuclear accumulation and transcriptional functions, thereby inhibiting ferroptosis and ferroptosis-mediated ICD. Low expression of NOLC1 could be a new biomarker for identifying GC patients who may benefit from Cis plus anti-PD-1 treatment and targeting NOLC1 could increase the treatment response of anti-PD-1 plus Cis.

## Materials and Methods

### Cell culture

HEK293T, MGC-803, and MKN-45 cell lines were purchased from the National Infrastructure of Cell Line Resource (Beijing, China). All cells were cultured in DMEM (GIBCO, USA) supplemented with 10% fetal bovine serum (FBS; Gibco, USA) and 1% anti-anti (100 U/ml; Gibco, USA). For Cisplatin resistant cell line construction, MGC-803 cells were treated with cisplatin (1 μM) for 5 month, then treated cells with cisplatin (5 μM) for 1month. All the cells were maintained at 37°C in a 5% CO_2_ cell culture incubator.

### RNA isolation and RT**□**qPCR

According to the manufaturer’s instructions, RNA was extracted from cells and tissues using TRIzol (Invitrogen, USA). Subsequently, using a reverse transcription kit (Takara, Dalian, China) transcribe RNA into cDNA. The expression levels of RNA transcripts were analyzed using a Bio-Rad CFX96 real-time PCR system (Bio-Rad, USA). All samples were normalized to GAPDH.

### Western blot

Collect cell samples and wash twice with cold PBS. After centrifugation for 5 min, lysed cells with RIPA lysis buffer (Beyotime, China) containing 1% PMSF and incubated on ice for 40 min. After centrifugation at 13,000 rpm for 20 minutes at 4 °C, the protein supernatant is collected. Proteins were fractionated by SDS–PAGE, transferred to PVDF membranes, blocked in 5-10% nonfat milk in TBS-Tween-20, and then treated with specific primary and secondary antibodies for 12h at 4°C. Finally, the blots were detected using enhanced chemiluminescence detection (NCM China). Intensities were analyzed using ImageJ software and relative expression was normalized to that of GAPDH.

### H&E and immunohistochemistry staining assay

The tumor and organ samples were collected and fixed in 4% paraformaldehyde solution for 24 h at 4°C. Then dehydrated, paraffin-embedded, sectioned, and stained with H&E.

For immunohistochemical staining analysis, sections were repaired in sodium citrate buffer and subsequently blocked with H_2_O_2_ for 30 min. Then sections were blocked with 5% normal goat serum, 0.1% Triton X-100, and 3% H_2_O_2_ for 30 min at room temperature and subsequently incubated with the appropriate primary antibodies overnight at 4°C. Finally, IHC staining was performed with horseradish peroxidase (HRP) conjugates using DAB detection.

### CCK-8 assay

MGC-803 or MKN-45 cells were seed in 96-well plates and incubated overnight. Then treated cells with indicating drugs for 24h. Afterward, adding 10% Cell Counting Kit-8 (CCK-8, Dojindo, Japan) to cells incubating them at 37°C for 1-3 h. The absorbance of each well was measured by a microplate reader (Tecan Switzerland) set at 450 nm. All experiments were performed in triplicate.

### Colony formation assay

For the colony formation assay, 1 × 10^3^ MGC-803 cells or 2 × 10^3^ MKN-45 cells are seeded into each well of six-well plates. And the medium was renewed every 3 days. The colonies were fixed with methanol after two weeks and then stained with 0.1% crystal violet (Sigma-Aldrich, USA) for 30 minutes and washed twice with PBS. The number of colonies with more than 50 cells was counted.

### Cell apoptosis assay

MGC-803 or MKN-45 cells were plated in 6-well plates and incubated overnight. Then treated with the indication treatment. Cell apoptosis was detected using an AnnexinV-APC apoptosis detection kit (Multi Science, China, Hangzhou) according to the manufacturer’s protocols. Briefly, the collected cells were washed twice with PBS and with binding buffer. Cells were resuspended in 500 μL binding buffer containing 5 μL of Annexin V-APC and 10 μL of 7-AAD. The analysis was then carried out using flow cytometry (Beckman Coulter, USA).

### LDH release assay

MGC-803 or MKN-45 cells were plated in 96-well plates (1 × 10^4^ cells well^−1^) and incubated overnight. Then treated with the indication treatment. Extracellular LDH was detected using an LDH assay kit (Dojindo, Japan) according to the manufacturer’s protocols.

### Immunofluorescence analysis

MGC-803, MKN-45, or HEK-293T cells were seeded in glass-bottom cell culture dishes (2 × 10^5^ cells well^-1^) and incubated for 12 h. Cells were then treated with PBS or cisplatin for 24 hours. Cells were then fixed with 4% paraformaldehyde for 20 min and blocked with 3% bovine serum albumin in PBS for 1 h. The fixed cells were then incubated with primary antibodies (NOLC1, p53, anti-HA tag, anti-Flag tag, HMGB1, γ-H2AX, and CRT) at 4 °C for 12 h. Subsequent incubation with an Alexa Fluor 562-conjugated anti-mouse or Alexa Fluor 647-conjugated anti-rabbit secondary antibody at room temperature for 2 h. DAPI was used to stain nuclear. Immunofluorescence images were captured using a confocal microscope (Nikon Japan). And analyzed by Image J software.

### FCM analysis of total ROS, mitoPeDPP, and mitochondrial membrane potential

MGC-803 and MKN-45 cells were treated with PBS or the indicated Cis concentration for 48 h, harvested cells and resuspended in 500 μL of PBS containing 1 μM DCFH-DA (Thermo Fisher, USA), 5 μM mitoPeDPP (Dojindo, Japan) or 1 μM JC-1 (Beyotime, China) and incubated at 37 °C for 30 minutes. The cells were analyzed by flow cytometer (Beckman Coulter, USA).

### Intracellular total ROS, MitoPeDPP, and mitochondrial membrane potential fluorescence imaging

Intracellular ROS were detected using a DCFH-DA probe (Thermo Fisher, USA). Havest cells and treated with DCFH-DA (1 μM) at 37 °C for 40 min. The fluorescence signals were detected by confocal microscopy (Nikon Japan). Intracellular mito-ROS were detected using the Mito-PeDPP probe. Harvest cells were isolated and treated with Mito-PeDPP (5 μM) for 1 hour at 37 °C. The fluorescence signals were detected by confocal microscopy (Nikon Japan). Mitochondrial membrane potential was detected with a JC-1 probe (Beyotime, China). The collected cells were treated with JC-1 (1 μM) for 20 min at 37 °C. The fluorescence signal was then detected by confocal microscopy (Nikon, Japan).

### H_2_O_2_ content assay

Total intracellular H_2_O_2_ was detected using a ROS-Glo™ H_2_O_2_ Assay Kit (Promega, USA). Briefly, MGC-803 or MKN-45 cells were plated at the desired density in ≤80 μL of medium in 96-well test plates with opaque walls. Treated cells with the indicated concentrations of cisplatin for 24 hours. Then 20 µl H_2_O_2_ substrate solution was added to the cell culture medium and incubated for 6 hours. Then add ROS-Glo™ Detection Solution and incubate for 20 minutes

### Transmission electron microscopy

MGC-803 cells were seeded in 6-well plates and treated with PBS or cisplatin for 24 h. Then the cells were collected and fixed with 3% glutaraldehyde in 0.1 M phosphate buffer, then fixed with OsO4. The cells were then dehydrated, and sectioned at 60–80 nm. Subsequently stain cells with uranyl acetate and lead nitrate. Finally, observation sections under a JEM-1230 transmission electron microscope (JEOL, Japan).

### Co-IP

Seeded HEK-293T cells in 6 cm cell culture dishes. When reached 70% density, the cells were transfected with the p53-HA, NOLC1-Flag, or Ub-MYC plasmid. Cells were collected 48 h later and lysed using IP lysis buffer (containing 1% protease inhibitor) for 30 min on ice. Then, anti-Flag magnetic beads were used to immunoprecipitate the NOLC1-Flag protein. The protein-antibody-magnetic bead complexes were dissociated by boiling using a metal bath for 10 min. The absorbed proteins were resolved by SDS□PAGE and subjected to immunoblotting with the indicated primary and secondary antibodies. The cell lysates were analyzed as an input.

### Chromatin immunoprecipitation (ChIP-Seq)

HEK293T cells were seeded in 6 cm cell culture dishes. When reached 70% density, the cells were transfected with p53-HA or NOLC1-Flag plasmids. The cells were harvested and processed with a SimpleChIP Enzymatic Chromatin IP Kit (CST, USA) following the manufacturer’s protocol. Briefly, after being fixed with 1% formaldehyde for 10 min at room temperature and lysed with Lysis Buffer-Protease Inhibitor solution, the cells were digested with micrococcal nuclease and immunoprecipitated with rabbit anti-HA antibodies. Then, DNA−protein complexes were eluted in extraction buffer, and incubated overnight at 65 °C to reverse the crosslinks. Finally, DNA was purified using spin columns for sequencing. DNA libraries and sequence was proceeded by Cosmos Wisdom Biotech Co., Ltd (Hangzhou, China)

### mRNA sequencing

Seeded MGC-803 and MGC-803-CR cells in 6 cm cell culture dishes, When reached 70% density, extracted by TRizol (ThermoFisher, USA). RNA intergrity was evaluated with a 1.0% agarose gel. Then RNA samples were send to Cosmos Wisdom Biotech Co., Ltd (Hangzhou, China) for sequencing.

### Luciferase reporter assay

HEK293T cells were seeded in 6 cm cell culture dishes. When reached 70% density, the cells were transfected with indicated reporters bearing an ORF encoding firefly luciferase, and pRL-Luc containing the Renilla luciferase ORF as the internal control for transfection. Briefly, luciferase assays were performed using a dual luciferase assay kit (Promega), luciferase activity was quantified with a microplate reader, and firefly luciferase activity was normalized to Renilla luciferase activity as the internal control.

### Animal experiments

The animal experiments procedures were approved by the Institutional Animal Care and Use Committee of Wenzhou Institute, University of Chinese Academy of Science (approval number: WIUCAS23112802). 4-week-old male BALB/c-nu mice were purchased from Vital River Laboratory Animal Technology Co., Ltd. (Shanghai). 4-week-old male 615 mice were purchased from ZiYuan Laboratory Animal Technology Co., Ltd. (Hangzhou). All mice were raised in specific-pathogen-free (SPF) animal rooms.

To establish the MGC-803 tumor-bearing model, a MGC-803 tumor-bearing model was established by subcutaneous injection of lentivirus (shNC or shNOLC1)-transfected MGC-803 cells (5 × 10^6^) into the right flanks of BALB/c male athymic nude mice. When the tumor volume reached about 80 mm^3^, the mice were randomly divided into two groups and intravenously administered PBS or cisplatin (40 μg/mouse) every 3 days for a total of 6 times.

To establish the MFC tumor-bearing model, an MFC tumor-bearing model was established by subcutaneous injection of lentivirus (shNC or shNOLC1)-transfected MFC cells (2 × 106) into the right flanks of 615 male mice. When the tumor volume reached about 100 mm3, randomly divided mice into four groups (n = 5) and intravenously administered PBS, PD-1 (100 μg/mouse), Cis (40 μg/mouse), or PD-1 combined with cisplatin every 3 days for a total of 3 times.

### Enzyme-Linked Immunosorbent Assay (ELISA)

Blood samples were collected and solidified at 4°C for 2 h. Then centrifuged at 2000 × g for 20 min at 4°C to obtain the blood serum. Cytokine and blood biochemistry levels were detected using ELISA kits according to the manufacturer’s protocol.

### In vivo Liperfluo analysis

Freshly frozen tissue was sectioned at a thickness of 5 µm, and then the slices were incubated with a Liperfluo fluorescent probe for 1 h at 37°C, then stain nuclear with Hoechst 33342. Finally observed slices by using confocal microscopy.

### Peripheral blood lymphocyte analysis

To analyze the peripheral blood lymphocytes, blood sample were collected to obtain lymphocytes. Blood was separated and filtered to obtain single lymphocytes suspension. After that the lymphocytes were and blocked by Mice TruStain FcX (Fc Receptor Blocking Solution, BioLegend) for 10 min. Then cells were stained with a LIVE/DEAD Fixable Violet Kit (Invitrogen) and subsequently stained with the following anti-mouse antibodies: CD3-PE, CD4-FITC, CD8-APC/Cyanine7 according to the manufacturer’s methods. Finally, cells were analyzed via a flow cytometer (Beckman Coulter, USA) and data analysis was conducted using FlowJo software (BD, USA).

### Statistics

All data are shown as mean ± standard deviation (SD), and Statistical analyses were conducted using GraphPad Prism 9 (GraphPad Software, San Diego, CA). Student’s t test was used to determine the significance of differences between two groups. The difference of *p < 0.05 was considered to indicate statistical significance, and **p < 0.01 and ***p < 0.001 were indicate high significance.

## Acknowledgements

We would like to thank all the participants involved in this study. We acknowledge the efforts of the doctor Cifeng Cai of the Cosmos Wisdom Biotech., Ltd (Hangzhou, China) for sequencing services.

## Ethics Approval

Studies involving human participants were viewed and approved by the Ethicss Committee in Clinical Research (ECCR) of the First Affiliated Hospital of Wenzhou medical University) Acceptance Number: KY2022-202

## Data Availability Statement

The data that support the findings of this study are available in the supplementary material of this article.

