## Supplemental figures and methods for "NOLC1 Suppresses Immuno-chemotherapy by Inhibiting p53-mediated Ferroptosis in Gastric Cancer"

**Supplementary figure and figure legend**

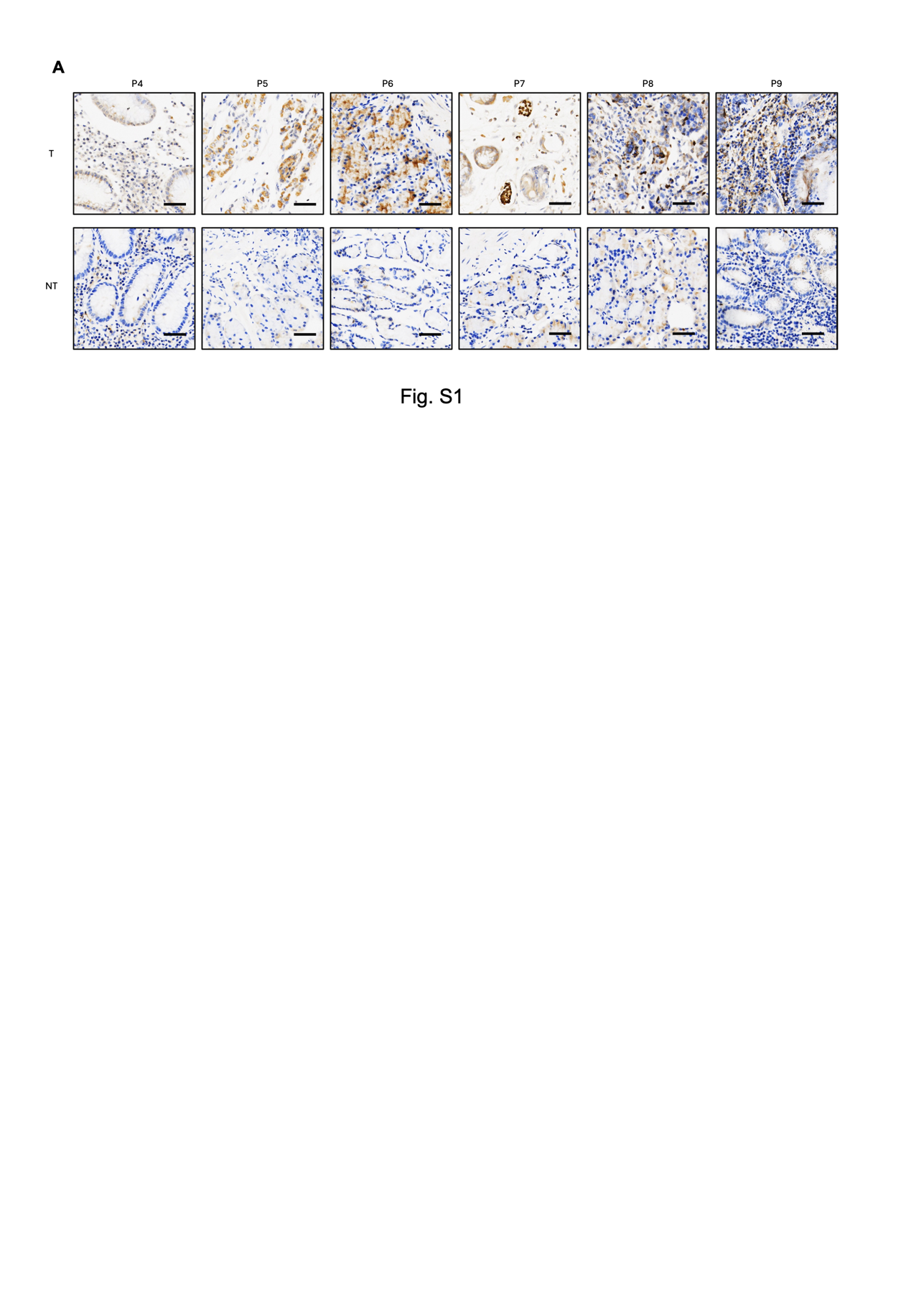

**Fig. S1 NOLC1 was highly expressed in GC tissues.**

**A** Immunohistochemistry (IHC) results of NOLC1 expression in gastric tumor (T) tissues and near tumor (NT) tissues.

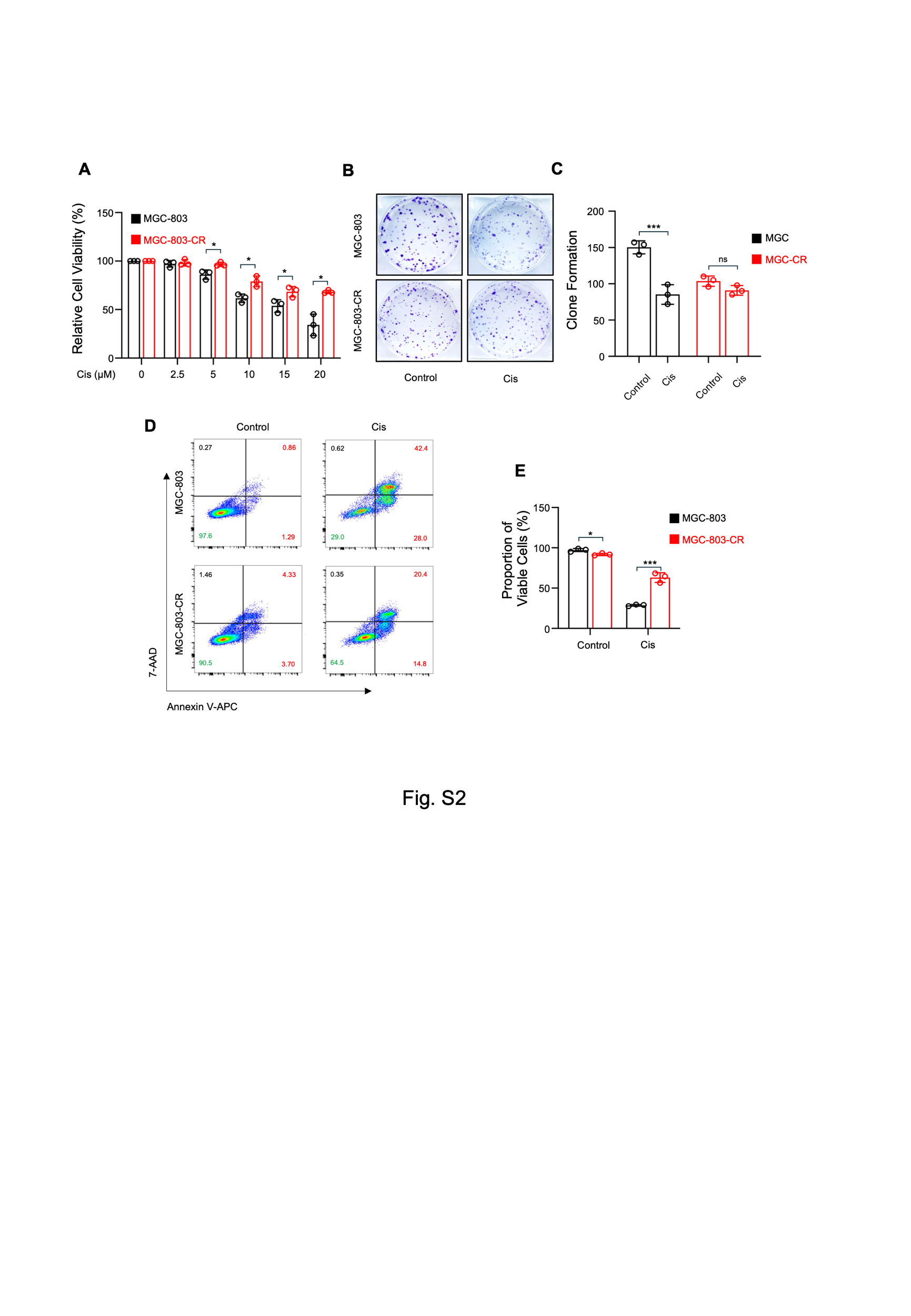

**Fig. S2 Cisplatin-resistant GC cell line was successfully constructed.**

**A** CCK-8 assay of MGC-803 and MGC-803-CR cells treated with different concentrations of Cis (*n* = 3).

**B, C** Colony formation assay of MGC-803 and MGC-803-CR cells treated with PBS or Cis (15 µM). (B) Representative images and (C) number of clones (*n* = 3).

**D, E** Annexin V-APC and 7-AAD staining of MGC-803 and MGC-803-CR cells treated with PBS or Cis (20 µM), as determined via FACS.

The data are presented as the means ± SDs. ns, nonsignificant; *p < 0.05; **p < 0.01; ***p < 0.001.

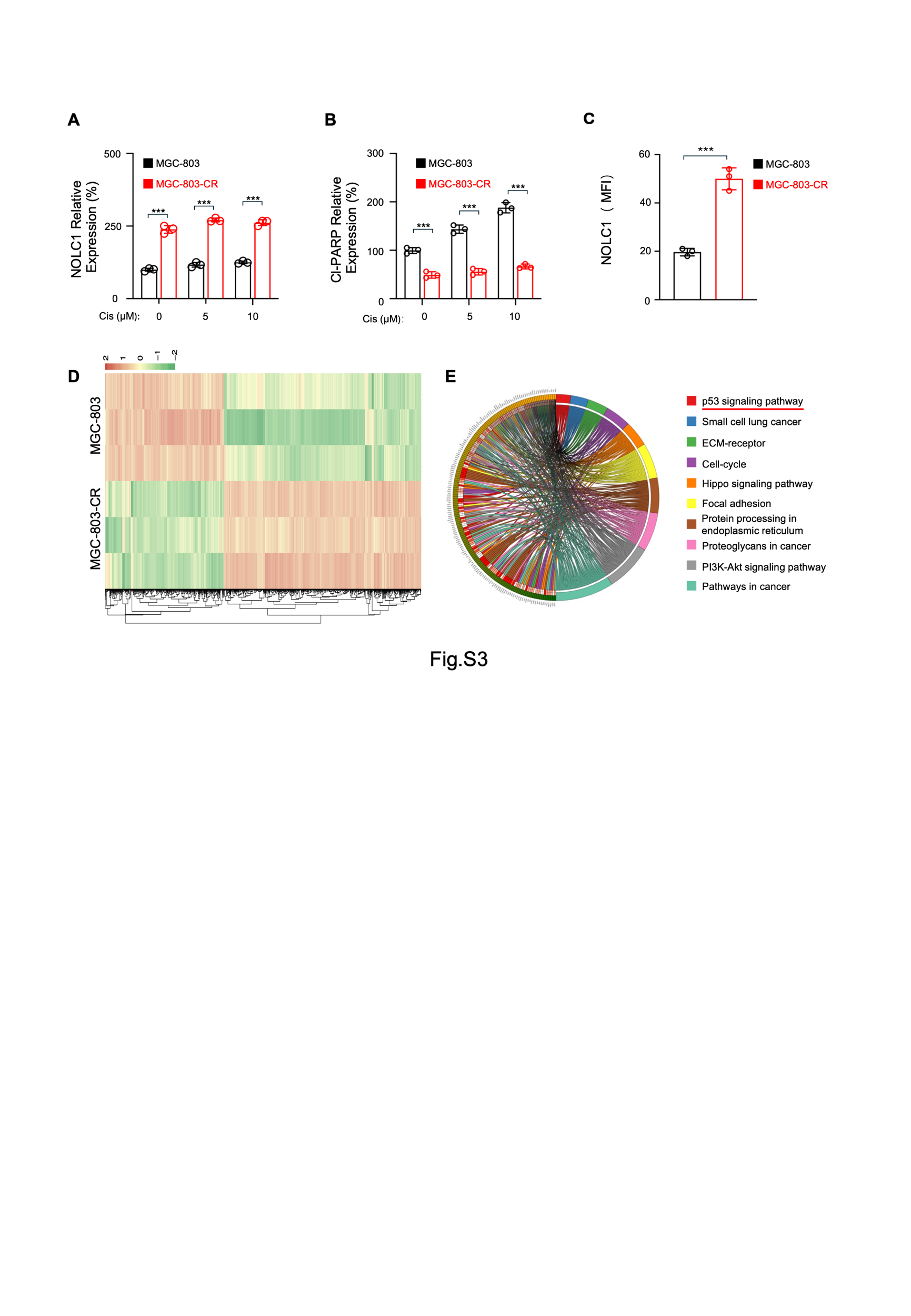

**Fig. S3** **NOLC1 was highly expressed in CR cells**

**A, B** Relative protein expression levels of (A) NOLC1 and (B) cleaved PARP (*n* = 3).

**C** Mean fluorescence intensity of NOLC1 (*n* = 3).

**D**. Heatmap of mRNA-seq data from MGC-803 and MGC-803-CR cells.

**E** KEGG analysis of the mRNA-seq data of MGC-803 and MGC-803-CR cells.

The data are presented as the means ± SDs. ***p < 0.001.

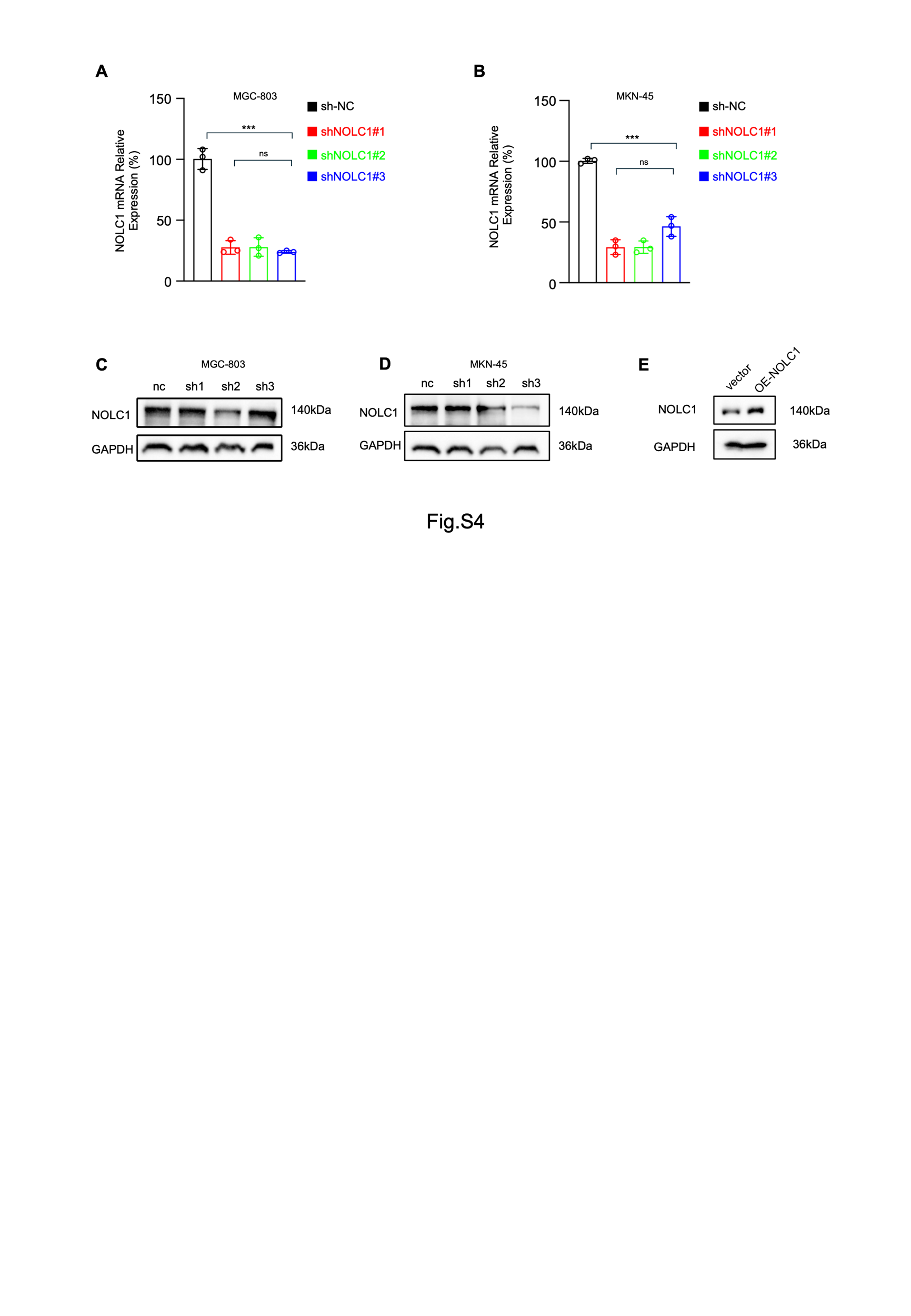

**Fig. S4 Knockdown or overexpression of NOLC1 in GC cell lines.**

**A, B** RT‒qPCR assay of NOLC1 mRNA levels in (A) MGC-803 cells and (B) MKN-45 cells (*n* = 3).

**C, D** Immunoblotting assay of NOLC1 in GC cells transduced with shNC or shNOLC1 lentivirus.

**E** Immunoblotting assay of NOLC1 in MGC-803 cells transduced with vector or NOLC1 plasmid.

The data are presented as the means ± SDs. ns, nonsignificant; ***p < 0.001.

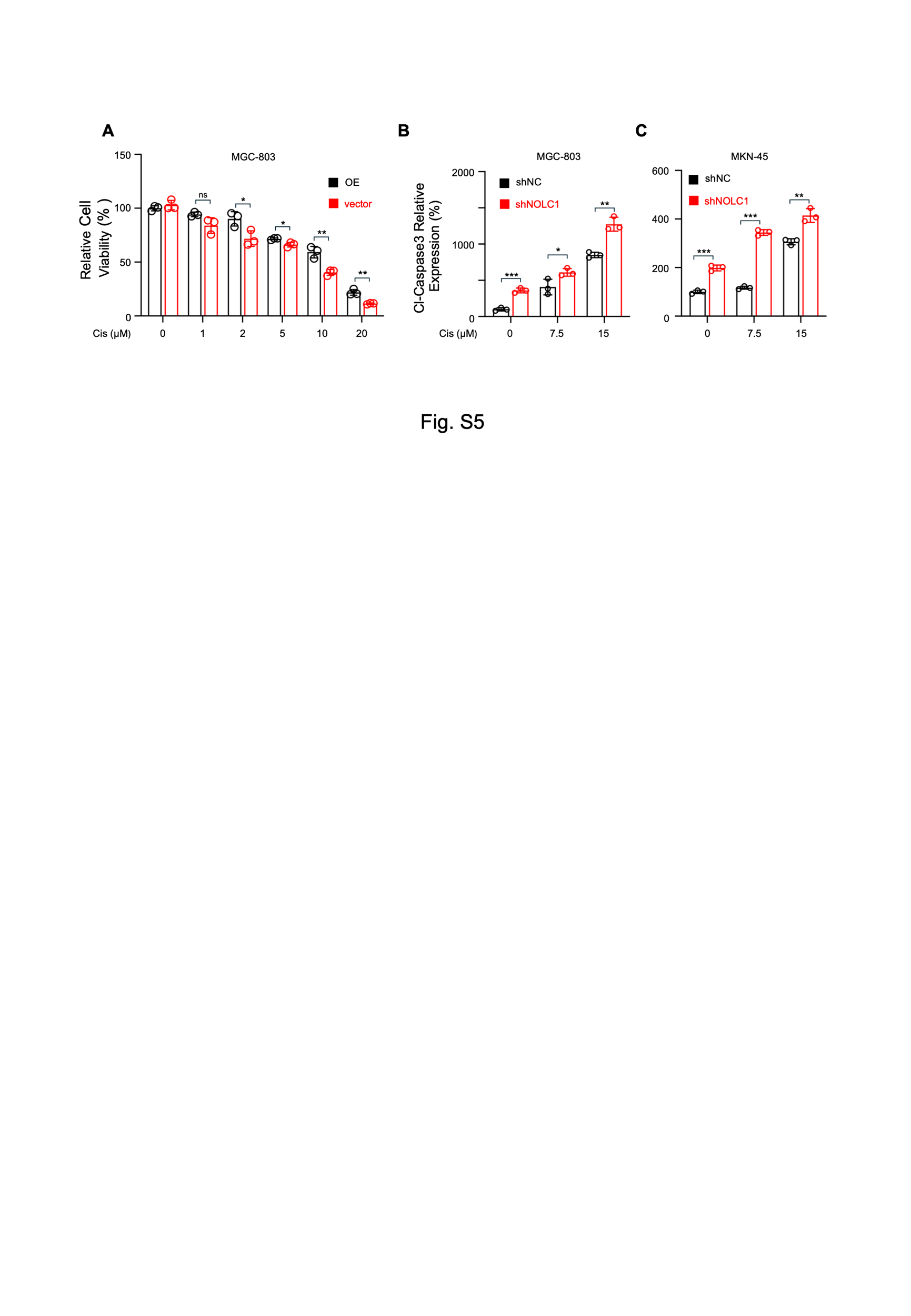

**Fig. S5 NOLC1 promoted Cis resistance in GC cells.**

**A** Viability of MGC-803 cells transduced with vector or NOLC1 plasmid and treated with the indicated concentrations of cisplatin (n = 3)

**B, C** Relative cleaved caspase3 expression level in GC cells transduced with shNC or shNOLC1 lentivirus and treated with indicated concentrations of Cis (*n* = 3): (A) MGC-803 cells and (B) MKN-45 cells.

The data are presented as the means ± SDs. ns, nonsignificant; *p < 0.05; **p < 0.01; ***p < 0.001.

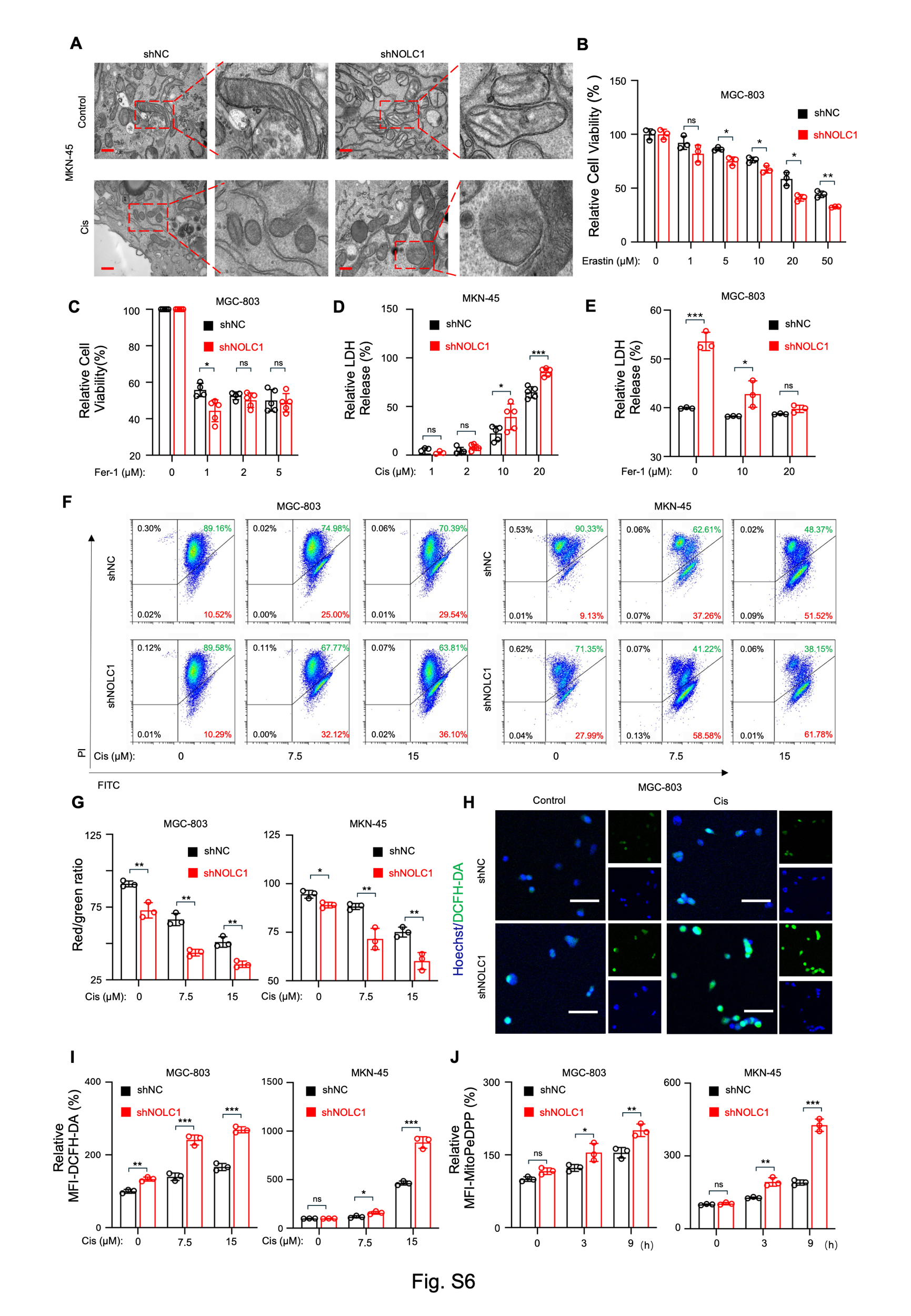

**Fig. S6** **NOLC1 deletion rendered GC cells susceptible to ferroptosis.**

**A** TEM image of MGC-803 cells transduced with shNC or shNOLC1 lentivirus and treated with PBS or Cis (15 µM); scale bar = 400 nm

**B** Viability of MGC-803 cells transduced with shNC or shNOLC1 lentivirus and treated with the indicated concentrations of erastin (*n* = 3).

**C** Viability of MGC-803 cells transduced with shNC or shNOLC1 lentivirus and treated with the indicated concentrations of Cis and Fer-1 (*n* = 5).

**D** LDH release analysis of MKN-45 cells transduced with shNC or shNOLC1 lentivirus and treated with different concentrations of Cis (*n* = 5).

**E** LDH release analysis of MGC-803 cells transduced with shNC or shNOLC1 lentivirus and treated with different concentrations of Fer-1 (*n* = 3).

**F** JC-1 fluorescence staining analysis of GC cells transduced with shNC lentivirus or shNOLC1 lentivirus and treated with indicated concentrations of Cis for via FACS.

**G** The ratio of red to green fluorescence of JC-1 in GC cells (*n* = 3).

**H** Representative DCFH-DA fluorescence images of MGC-803 cells transduced with shNC or shNOLC1 lentivirus and treated with indicated concentrations of Cis; scale bar = 75 µm.

**I** Relative mean fluorescence intensity (MFI) of DCFH-DA in GC cells transduced with shNC or shNOLC1 lentivirus and treated with indicated concentrations of Cis (*n* = 3).

**G** Relative mean fluorescence intensity (MFI) of MitoPeDPP in GC cells transduced with shNC or shNOLC1 lentivirus and treated with Cis (15 µM) for different time (*n* = 3).

**I** IHC staining analysis of GPX4 in MGC-803 tumor tissues after the indicated treatments; scale bar = 50 µm.

The data are presented as the means ± SDs. ns, nonsignificant; *p < 0.05; **p < 0.01; ***p < 0.001.

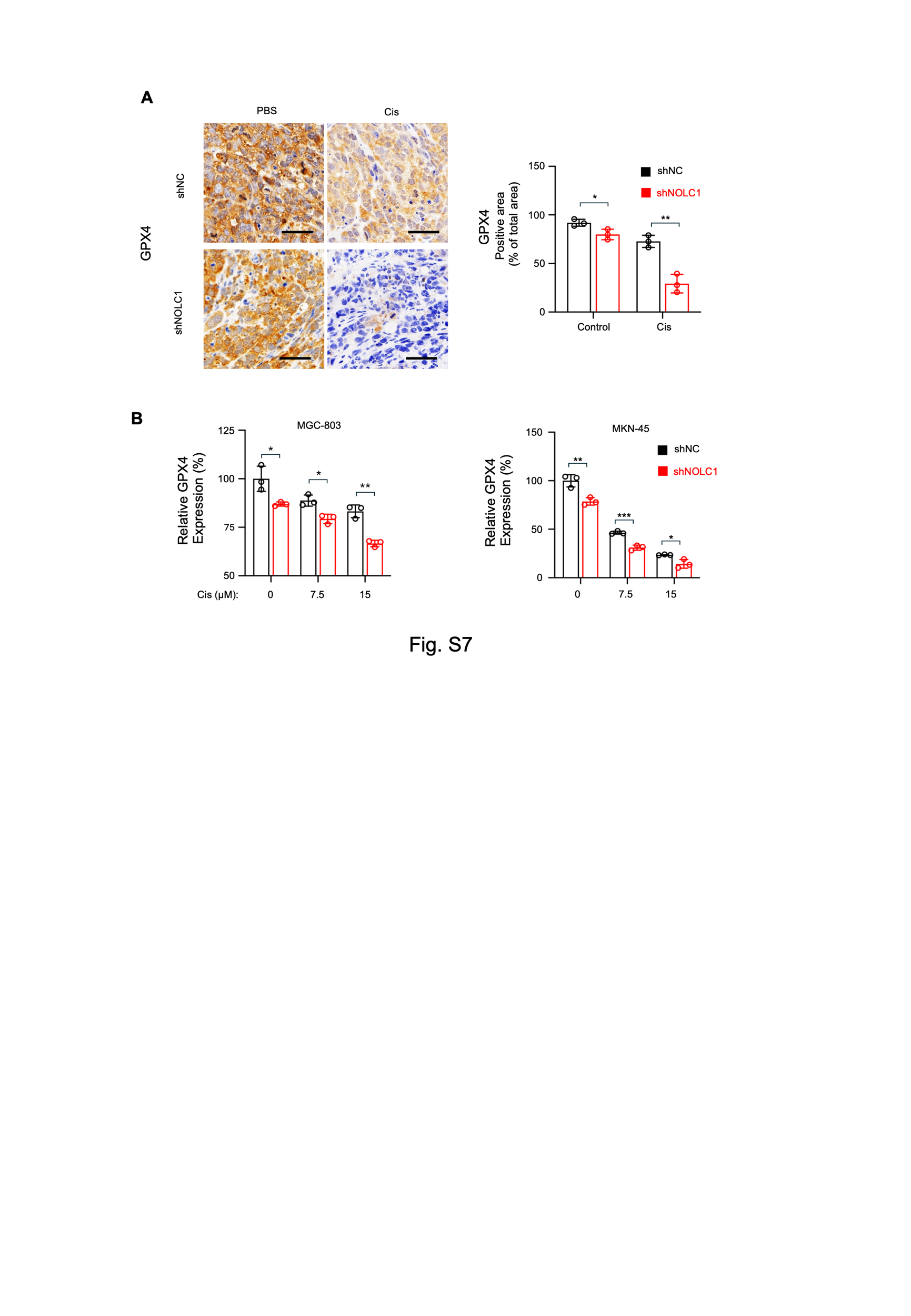

**Fig. S7 NOLC1 deletion rendered GC cells susceptible to ferroptosis.**

**A** IHC staining analysis of GPX4 in MGC-803 tumor tissues after the indicated treatments; scale bar = 50 µm.

**B** Relative protein expression of GPX4 in GC cells transduced with shNC or shNOLC1 lentivirus and treated with different concentrations of Cis (*n* = 3);

The data are presented as the means ± SDs. *p < 0.05; **p < 0.01; ***p < 0.001.

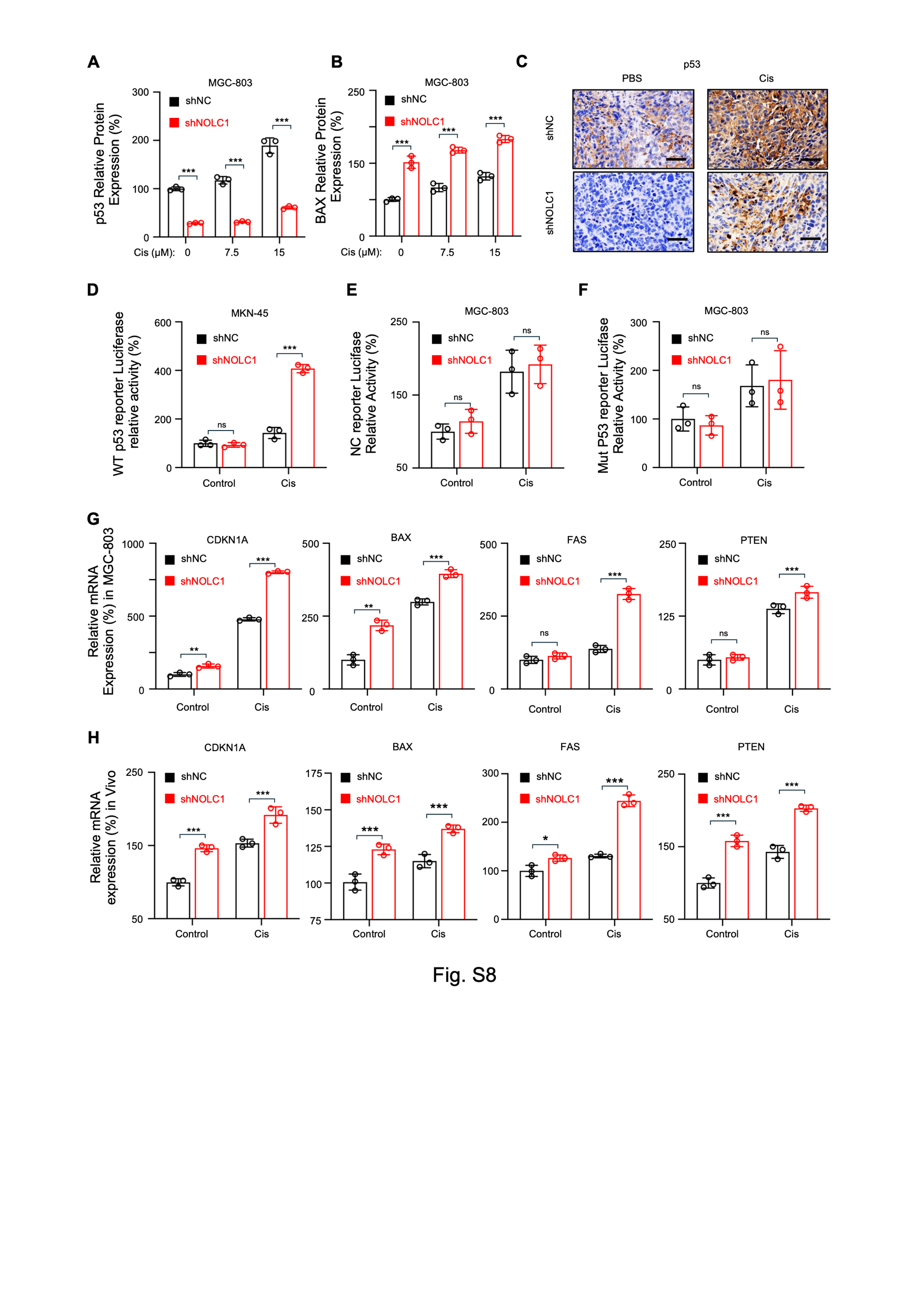

**Fig. S****8 NOLC1 inhibited p53 transcriptional activity.**

**A, B** Relative protein expression levels of (A) p53 and (B) BAX in MGC-803 cells transduced with shNC or shNOLC1 lentivirus and treated with different concentrations of Cis (*n* = 3).

**C** IHC staining of p53 in MGC-803 tumor tissues after different treatments; scale bar = 50 µm.

**D** Luciferase assay of WT p53 reporter transcription activity in MKN-45 cells transduced with shNC or shNOLC1 lentivirus and treated with PBS or Cis (15 µM) (*n* = 3).

**E** Luciferase assay of NC reporter transcription activity in HEK-293T cells transduced with shNC or shNOLC1 lentivirus and treated with PBS or Cis (15 µM) (*n* = 3).

**F** Luciferase assay of Mut p53 reporter transcription activity in HEK-293T cells transduced with shNC or shNOLC1 lentivirus and treated with PBS or Cis (15 µM) (*n* = 3).

**G, H** RT‒qPCR analysis of CDKN1A, BAX, FAS, and PTEN mRNA levels (G) in MGC-803 cells transduced with shNC or shNOLC1 lentivirus and treated with Cis (0 or 15 µM) (*n* = 3) and (H) in MGC-803 tumor tissues after the indicated treatment (*n* = 3).

The data are presented as the means ± SDs. ns, nonsignificant; *p < 0.05; **p < 0.01; ***p < 0.001.

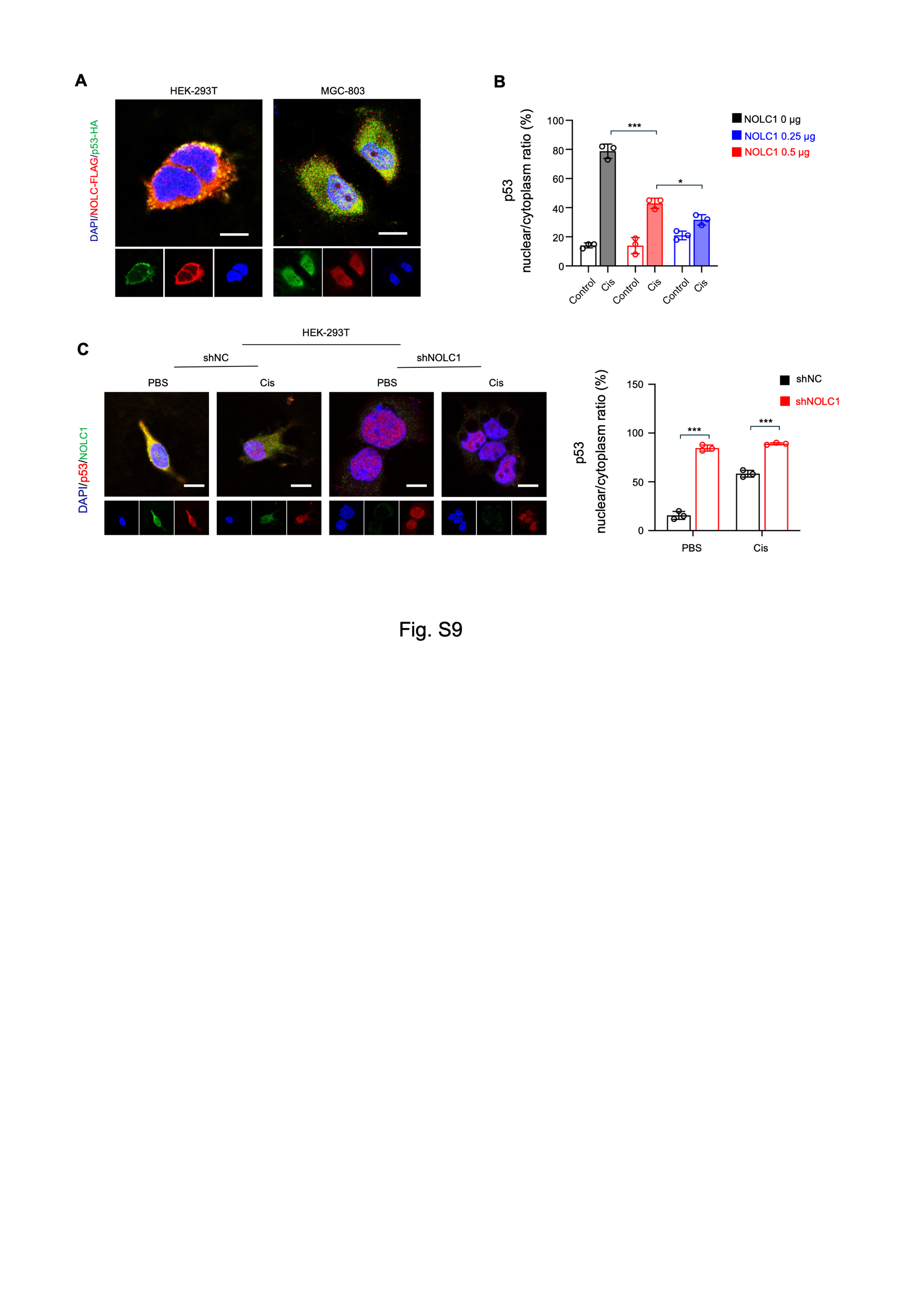

**Fig. S9** **NOLC1 combined with p53 and inhibited p53 nuclear accumulation.**

**A** Representative NOLC1 and p53 fluorescence images of HEK-293T and MGC-803 cells transfected with NOLC1-Flag and p53-HA plasmids; scale bar = 5 µm or 10 µm respectively.

**B** The nuclear/cytoplasm ratio of p53 in HEK-293T cells transfected with different amount NOLC plasmid.

**C** Representative immunofluorescence images of p53 and NOLC1 stained HEK-293T cells transduced with shNC or shNOLC1 lentivirus and treated with PBS or Cis (15µM); scale bar = 10 µm.

The data are presented as the means ± SDs. *p < 0.05; ***p < 0.001.

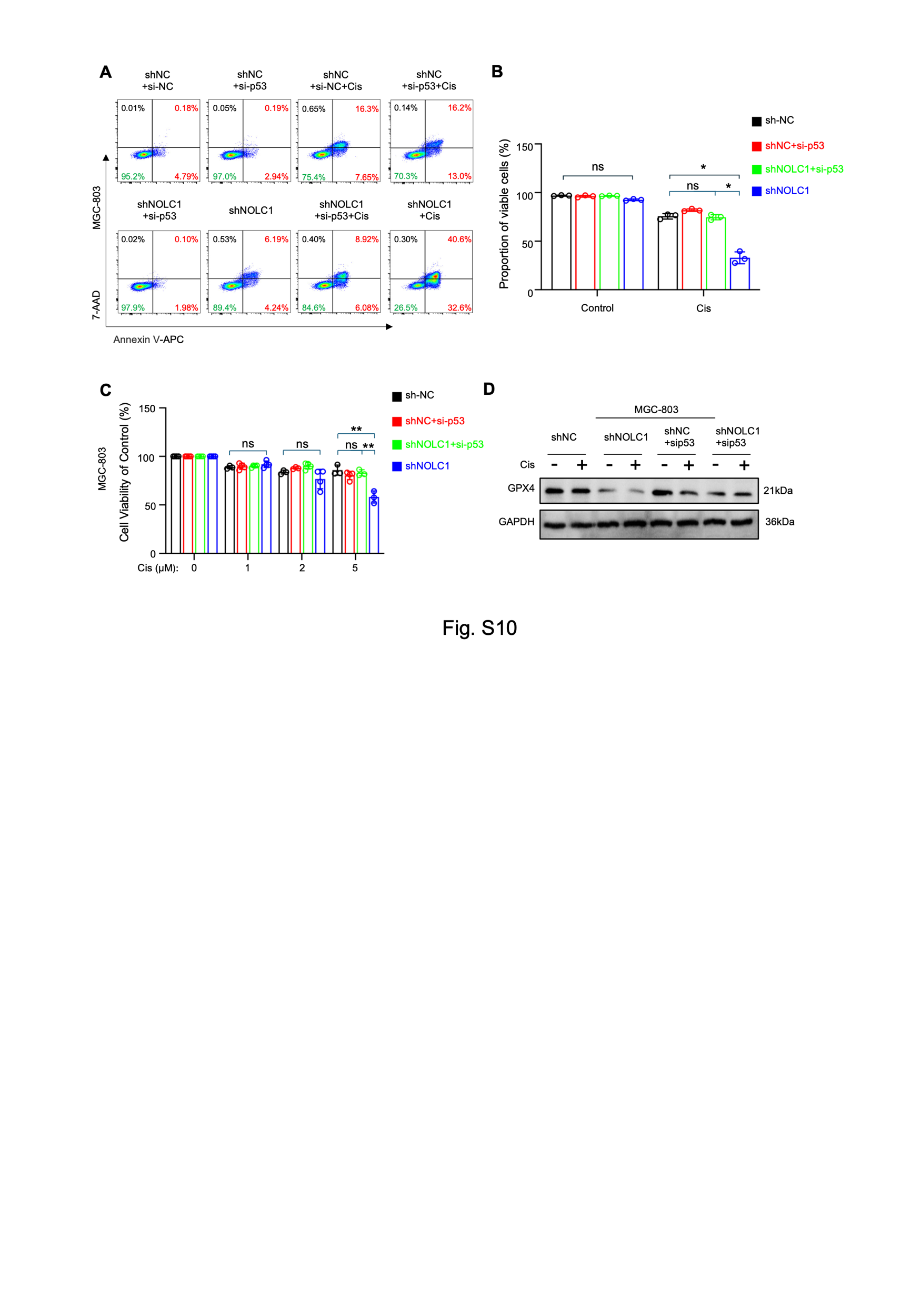

**F****ig. S10** **NOLC1 promoted Cis resistance by blocking p53 function in GC.**

**A, B** Annexin V-APC and 7-AAD staining of GC cells transduced with shNC or shNOLC1 lentivirus and transfected with si-NC or si-p53 siRNA and treated with PBS or Cis (15 µM), as analyzed via FACS.

**C** Viability of MGC-803 cells transduced with shNC or shNOLC1 lentivirus and transfected with si-NC or si-p53 siRNA and treated with different concentrations of Cis.

**D** Immunoblotting analysis of GPX4 protein levels in MGC-803 cells transduced with shNC or shNOLC1 lentivirus and transfected with si-NC or si-p53 siRNA and treated with 0 or 10 µM Cis.

The data are presented as the means ± SDs. ns, nonsignificant; *p < 0.05; **p < 0.01.

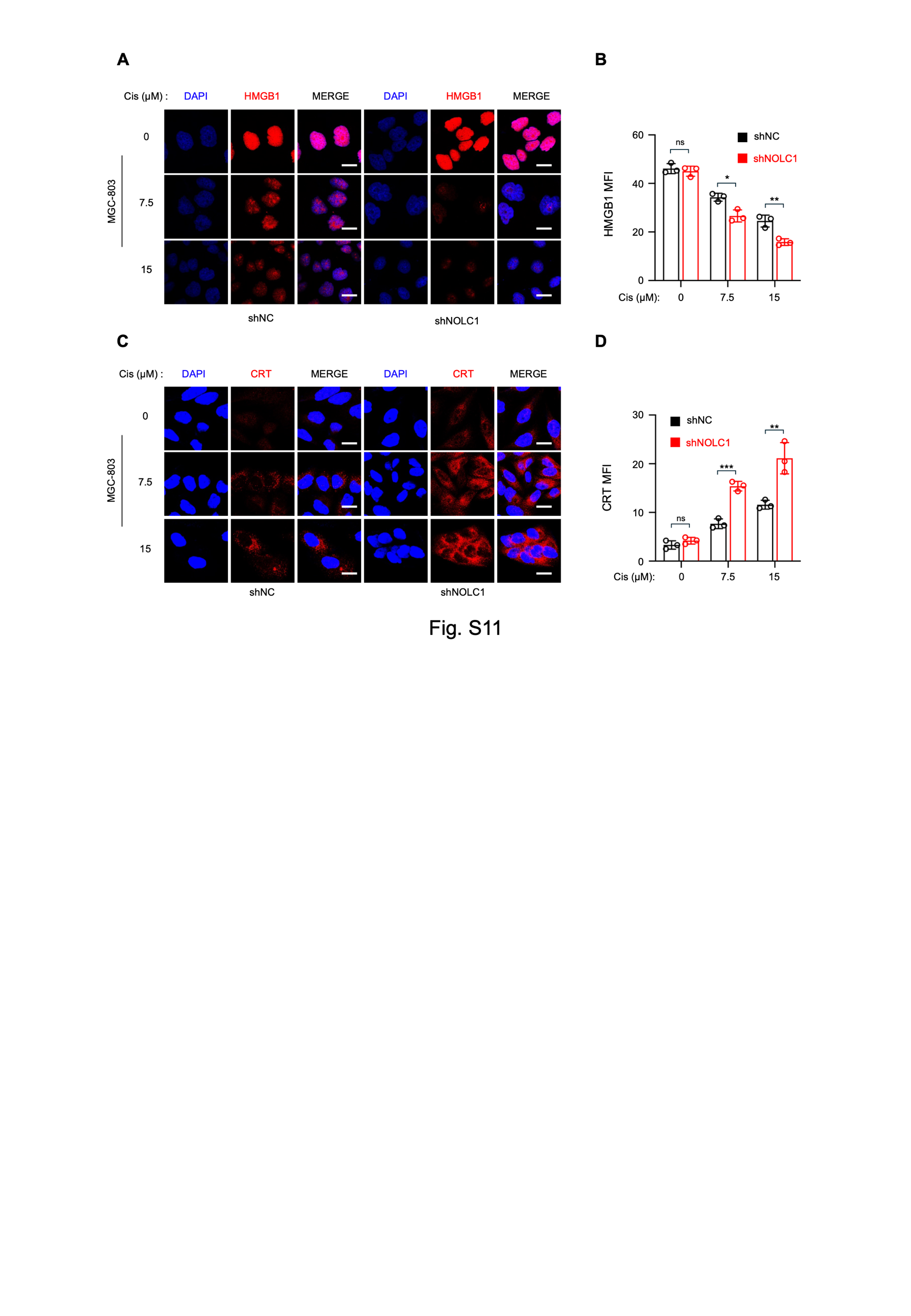

**Fig. S11** **NOLC1 knockdown increased the release of DAMPs induced by ferroptosis.**

**A, B** HMGB1 immunofluorescence staining of MGC-803 cells transduced with shNC or shNOLC1 lentivirus and treated with different concentrations of Cis. (A) Representative images; scale bar = 10 µm. (B) Mean fluorescence intensity of HMGB1 (*n* = 3).

**C, D** CRT immunofluorescence staining of MGC-803 cells transduced with shNC or shNOLC1 lentivirus and treated with different concentrations of Cis. (C) Representative images; scale bar = 10 µm. (D) Mean fluorescence intensity of CRT (*n* = 3).

The data are presented as the means ± SDs. ns, nonsignificant; *p < 0.05; **p < 0.01; ***p < 0.001.

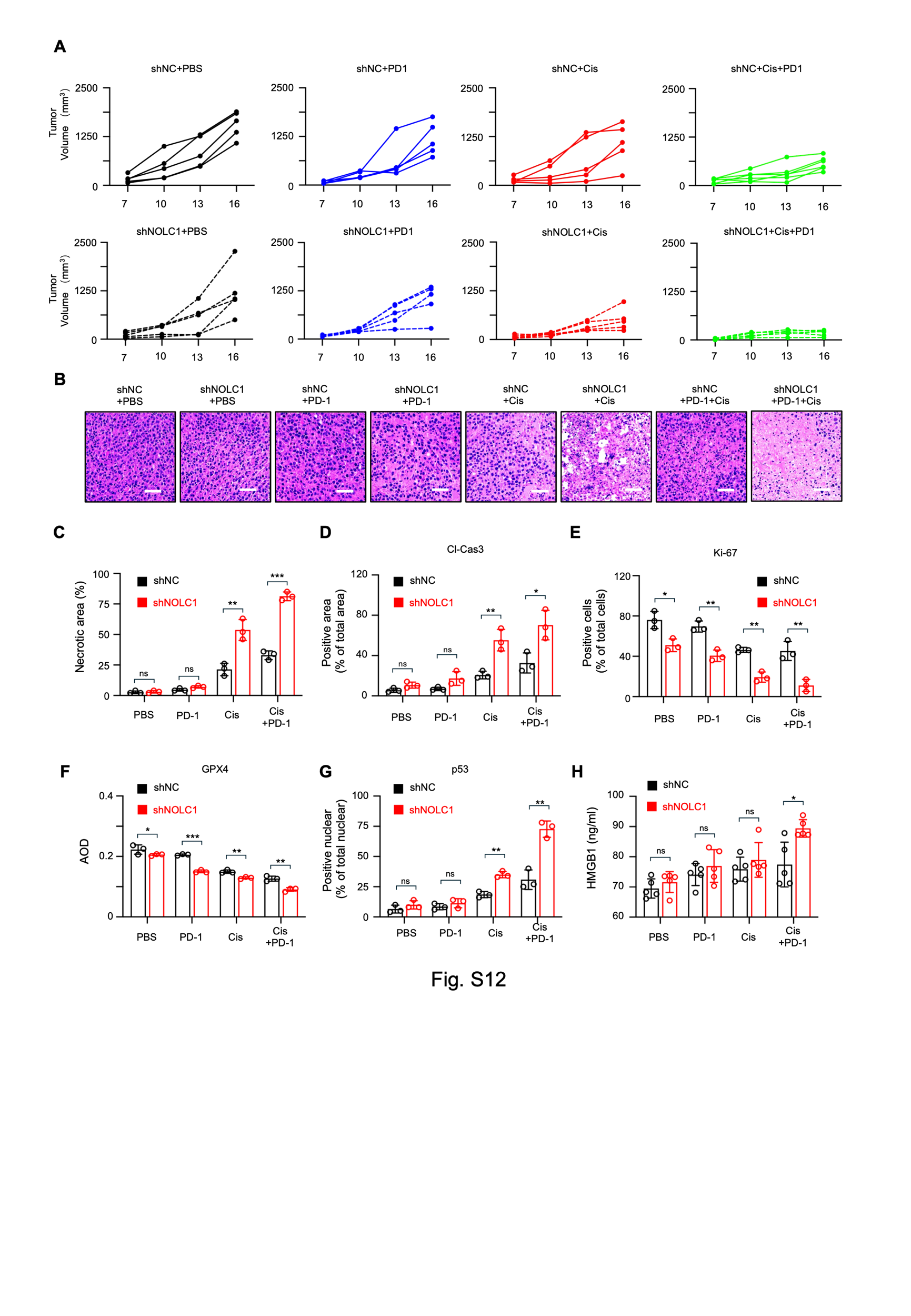

**Fig. S12** **NOLC1 knockdown increased the combination treatment efficacy of anti-PD-1 plus Cis.**

**A** Growth curves of MFC tumors transduced with shNC or shNOLC1 lentivirus and subjected to different treatments.

**B** H&E staining of MFC tumors transduced with shNC or shNOLC1 lentivirus after the indicated treatments.

**C** Quantification of the necrotic area in the tumor tissue of the mice given the different treatment (*n* = 3).

**D**-**G** Quantification of protein expression in tumor tissue from mice given the indicated treatment (*n* = 3). (D) cleaved caspase-3, (E) Ki-67, (F) GPX4, and (G) p53 nuclear positive ratio.

**H** The serum levels of HMGB1 (*n* = 5).

The data are presented as the means ± SDs. ns, nonsignificant; *p < 0.05; **p < 0.01; ***p < 0.001.

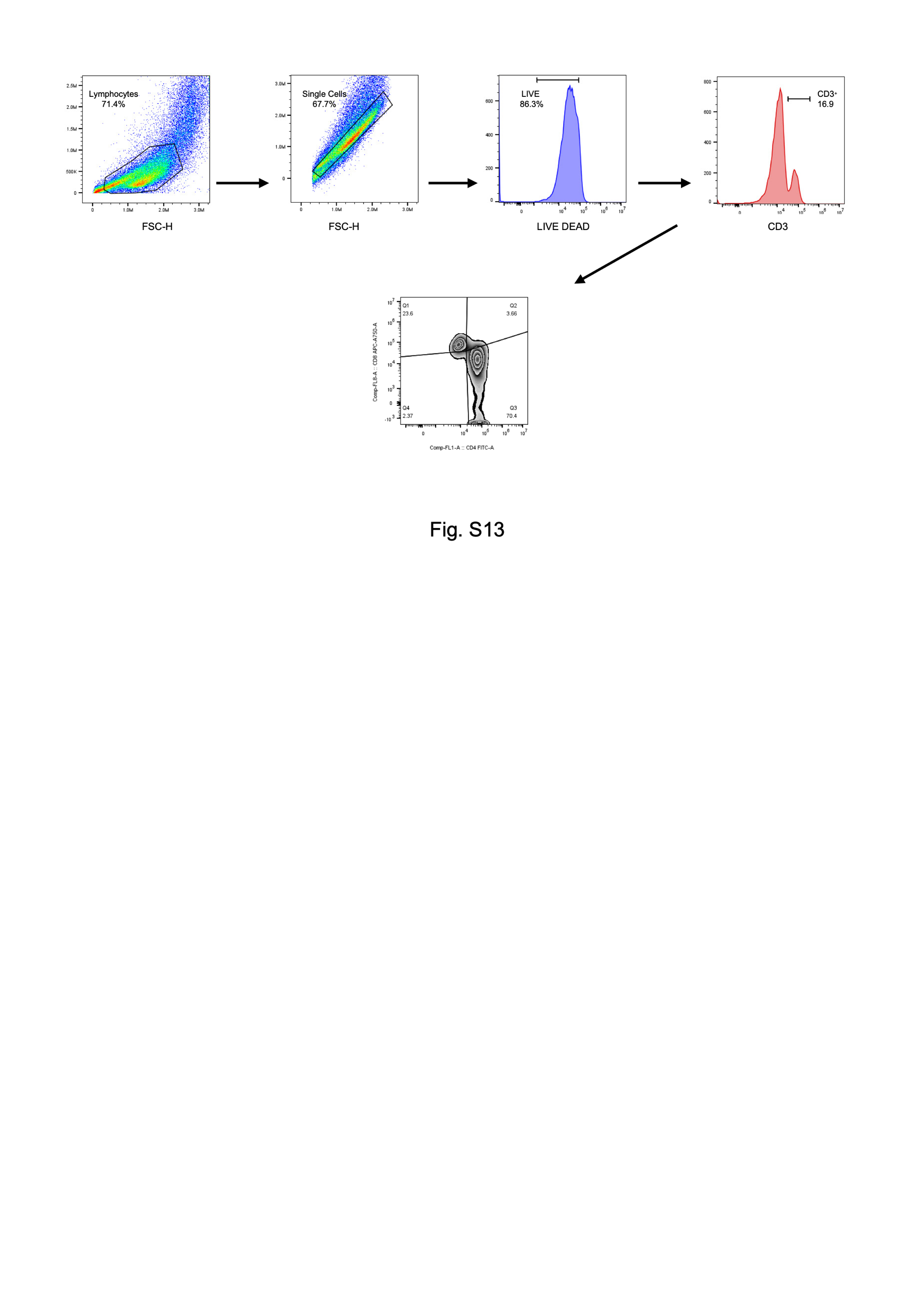

**Fig. S13 Gating strategy of flow cytometry for peripheral blood lymphocyte analysis**

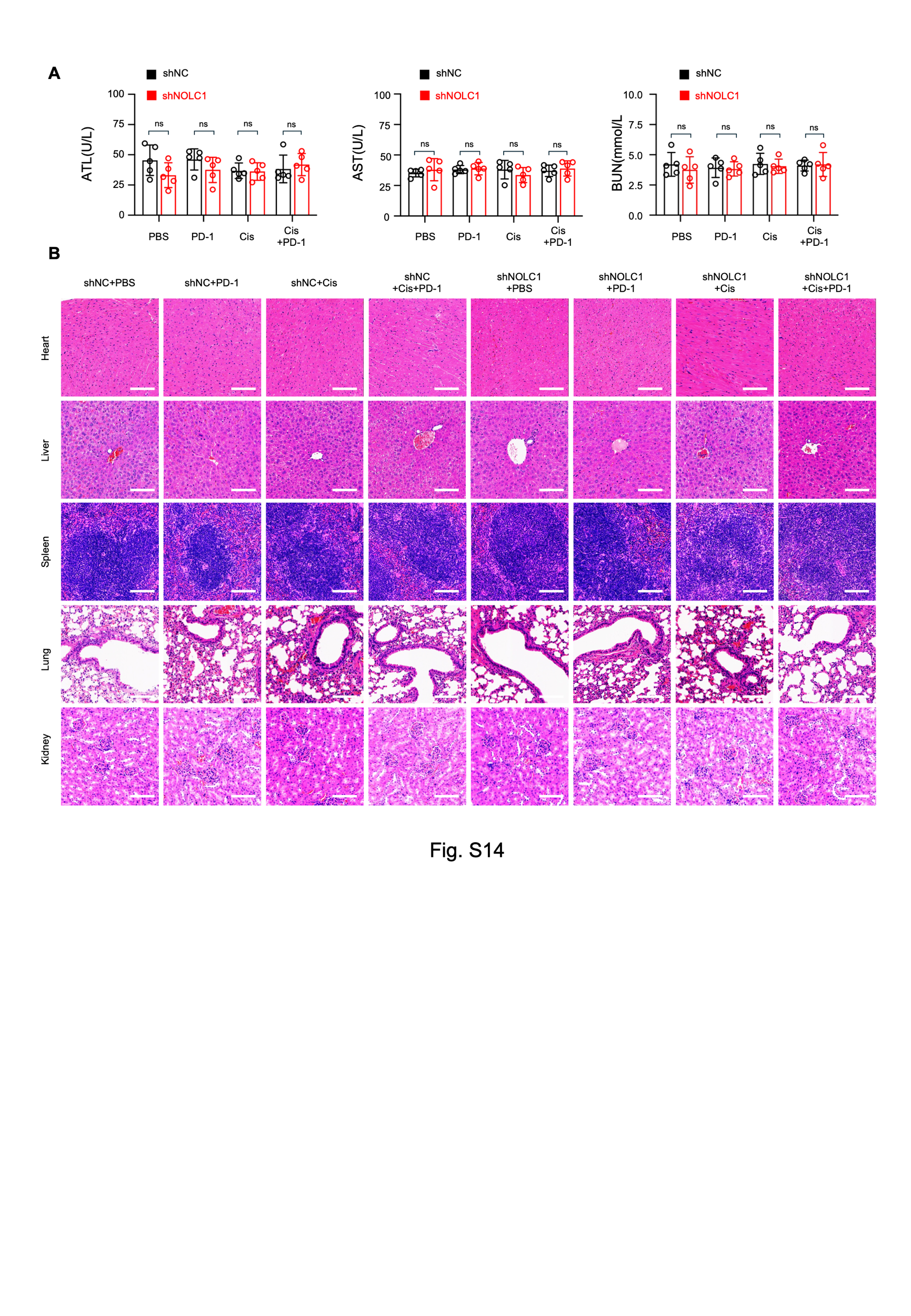

**Fig. S14 Biosafety of anti-PD-1 and Cis combination therapy.**

**A** Biochemical analysis of the serum of mice subjected to various treatments (*n* = 5).

**B** H&E staining of mouse heart, liver, spleen, lung, and kidney samples from different treatment groups. Scale bar = 100 μm.

The data are presented as the means ± SDs. ns, nonsignificant.

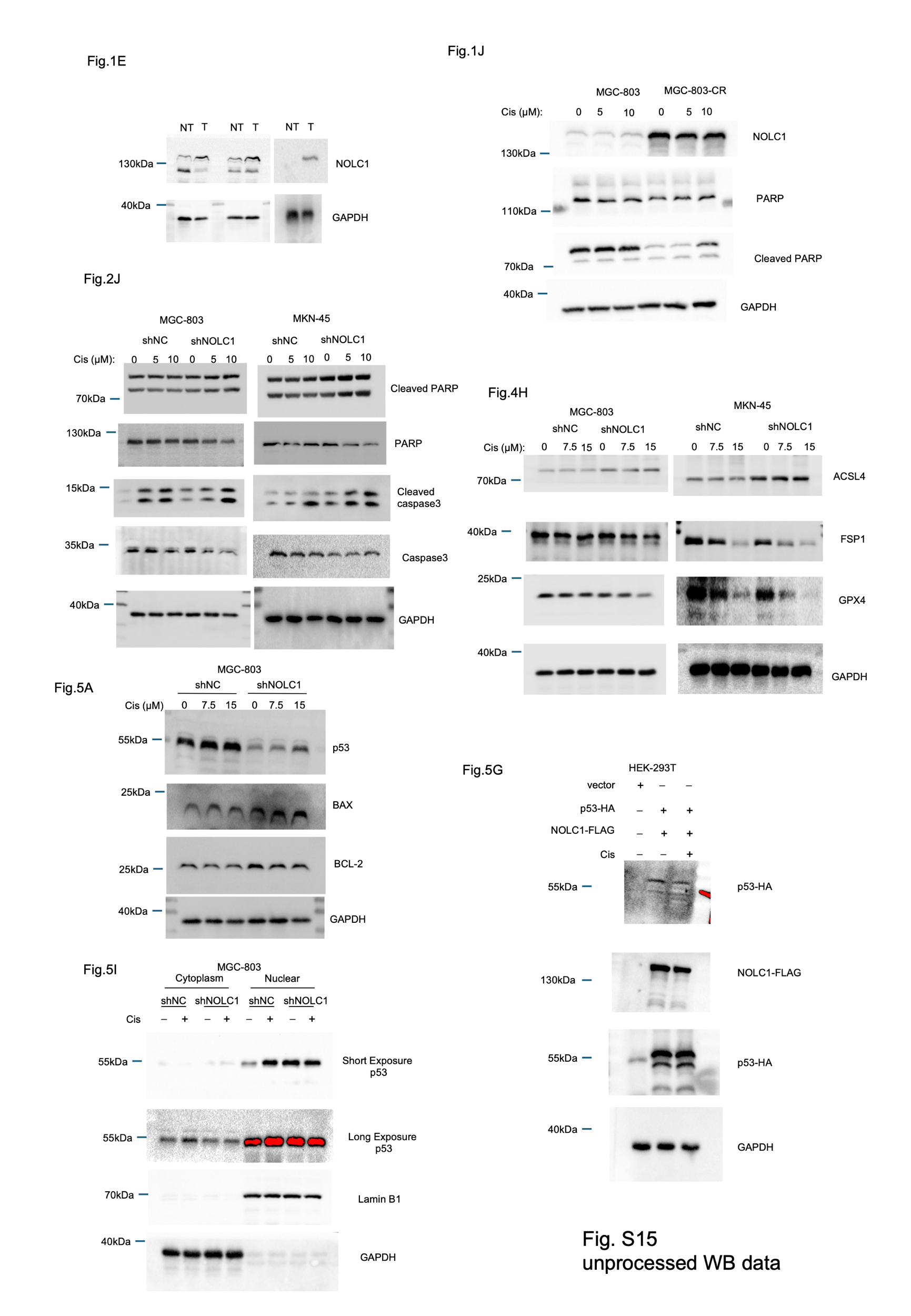

**Fig. S15 Unprocessed blots**

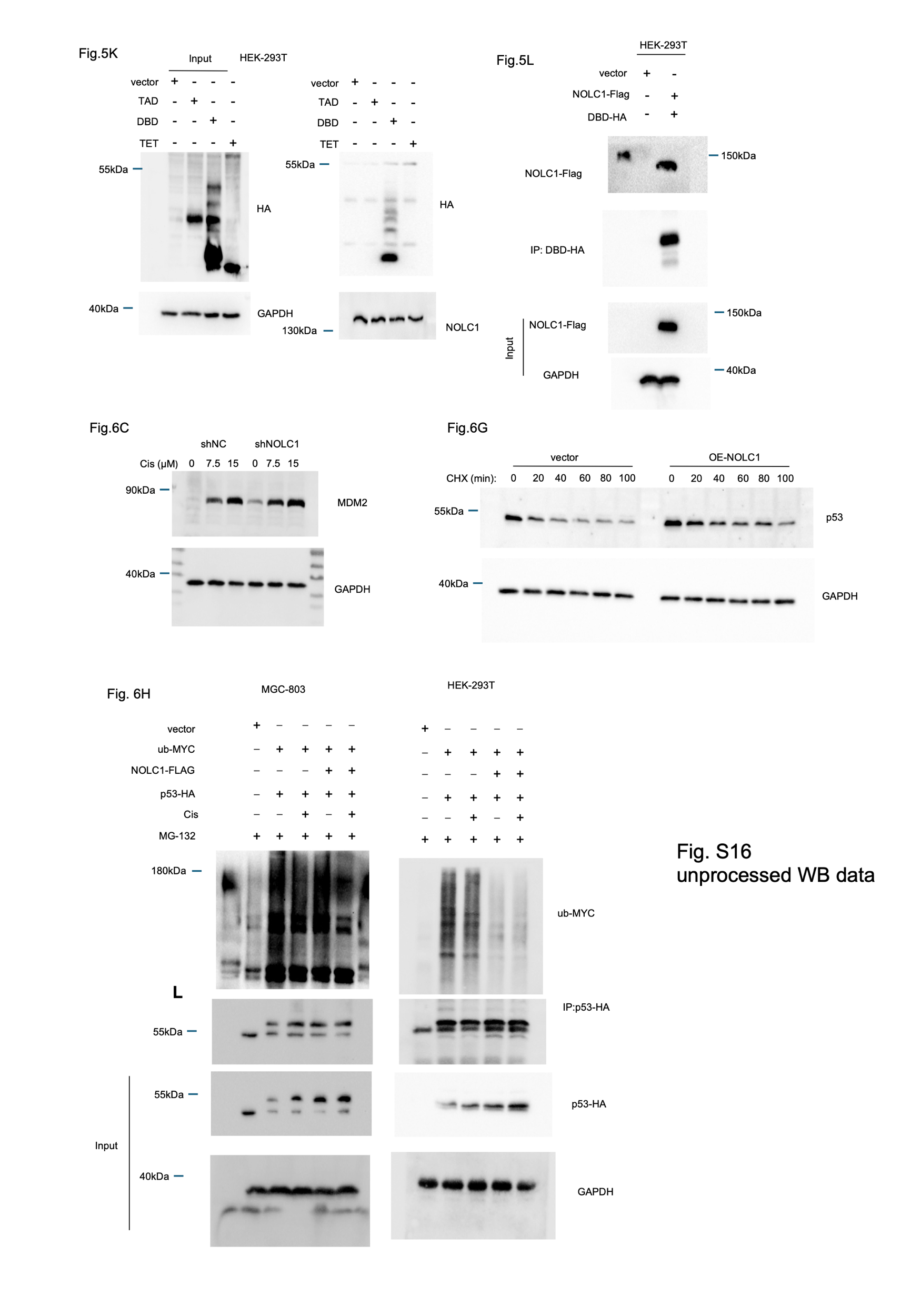

**Fig. S16 Unprocessed blots**

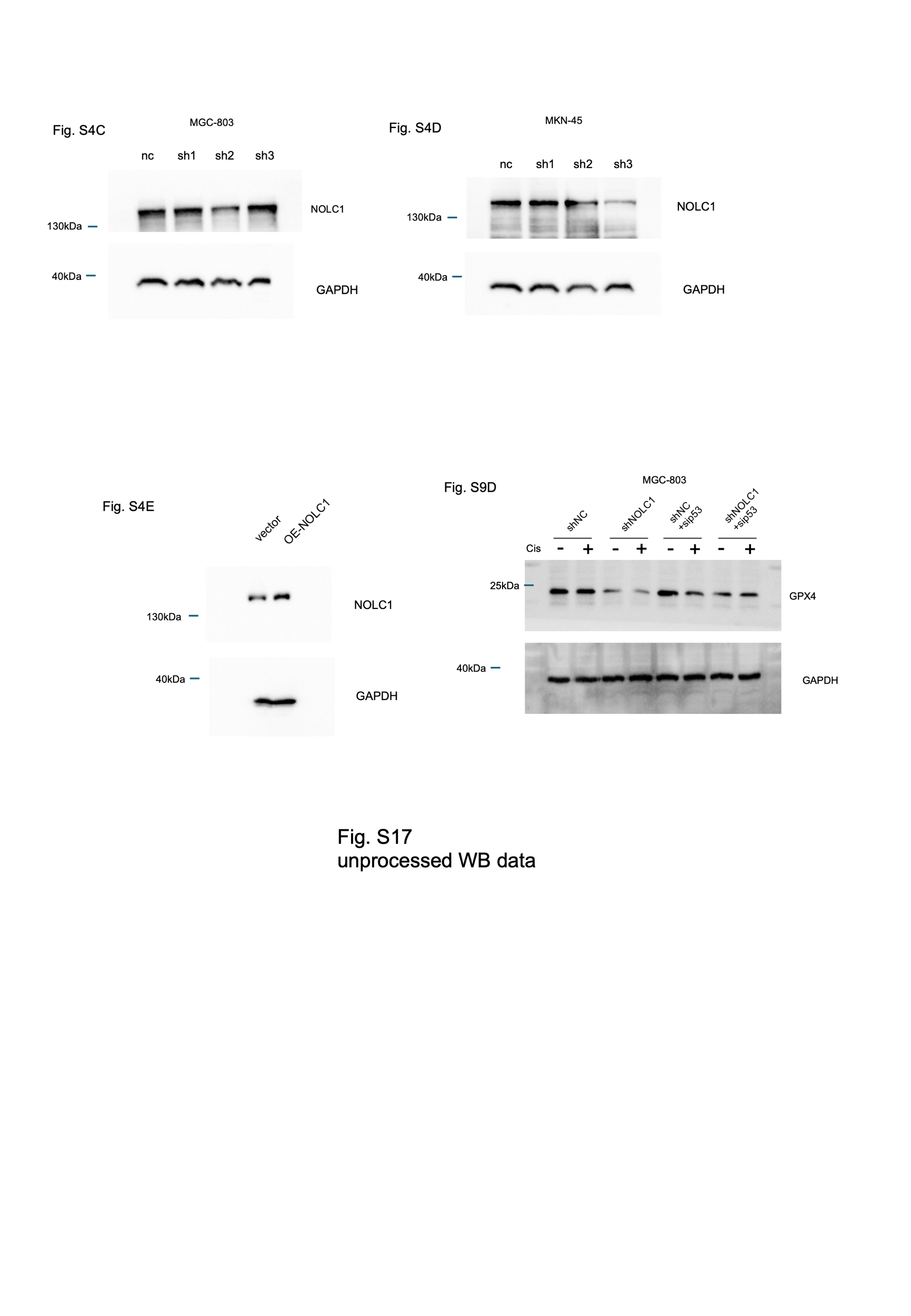

**Fig. S17 Unprocessed blots**

**Supplemental Materials and Methods**

**Expression plasmids**

Expression plasmids encoding Flag, HA- or MYC-tagged human NOLC1, p53, and Ub, in addition to reporter vectors containing the p53 promoter, were purchased from RiboBio. Lentivirus-NOLC1 was obtained from Taitool. All coding sequences were verified by DNA sequencing.

**Chemicals**

Ferrostatin-1, cisplatin, MG-132, and CHX were obtained from MCE.

**Antibodies**

NOLC1 (Cat: DF4264, RRID:AB_2836615), PARP (Cat: DF7198, RRID:AB_2839150), cleaved-PARP (Asp214) (Cat: AF7023, RRID:AB_2835327), P21 (Cat: AF6290, RRID:AB_2827699), FAS (Cat: AF5342, RRID:AB_2837827), MDM2 (Cat: AF0208, RRID:AB_2833395), and ACSL4 (Cat: DF12141, RRID:AB_2844946) antibodies were obtained from Affinity. Lamin B1 antibody (Cat. 12987-1-AP, RRID:AB_2136290), MYC tag antibody (Cat. 16286-1-AP, RRID:AB_11182162), p53 antibody (Cat. 60283-2-Ig, RRID:AB_2881401), caspase3/p17/p19 antibody (Cat. 19677-1-AP, RRID:AB_10733244), GAPDH antibody (Cat. 10494-1-AP, RRID:AB_2263076), PTEN antibody (Cat. 22034-1-AP, RRID:AB_2878977), DYKDDDDK tag antibody (Cat. 66008-4-Ig, RRID:AB_2918475), and HA-tag antibody (Cat. 51064-2-AP, RRID:AB_11042321) were obtained from Proteintech. Bcl-2 antibody (Cat: ab1828585), CD3 epsilon antibody (Cat: ab16669, RRID:AB_443425), and CD8 alpha antibody (Cat: ab217344, RRID:AB_2890649) were obtained from Abcam. Phospho-histone H2A.X (Ser139) antibody (Cat: 2577, RRID:AB_2118010), GPX4 antibody (Cat: 52455, RRID:AB_2924984), Ki-67 antibody (Cat: 9129, RRID:AB_2687446), and FSP1 antibody (Cat: 24972, RRID:AB_3090192) were obtained from CST. The donkey anti-mouse IgG (H+L) highly cross-adsorbed secondary antibody Alexa Fluor 647 (Cat: A31571, RRID:AB_162542), goat anti-rabbit IgG (H+L) cross-adsorbed secondary antibody Alexa Fluor 555 (Cat: A21428, RRID:AB_141784), goat anti-rabbit IgG (H+L) secondary antibody, HRP (Cat: 31460, RRID:AB_228341), and goat anti-mouse IgG (H+L) secondary antibody HRP (Cat: 31430, RRID:AB_228307) were obtained from Invitrogen. The InVivoMAb anti-mouse PD-1 antibody (Cat: BE0146, RRID:AB_10949053) was obtained from Bio X Cell.

**Kits**

Cell Counting Kit-8 (CCK-8) and a Cytotoxicity LDH Assay Kit (Cat: CK12) were obtained from Dojindo Laboratories. A dual-luciferase kit and H_2_O_2_ assay kit were obtained from Promega. Enzyme-linked immunosorbent assay (ELISA) kits for HMGB-1, IL-6, TNF-𝛼, IFN-𝛾, ALT, AST, and BUN were obtained from Jiangsu Meimian Industrial Co., Ltd. mRNA reverse transcription kits were obtained from Takara.

**Probes**

The DCFH-DA probe was obtained from Thermo Fisher Scientific. JC-1 probes were obtained from Beyotime Biotechnology. MitoPeDPP and LiperFluo probes were obtained from Dojindo Laboratories.

**Primes**

| NOLC1-F | TTCCTGCGCGATAACCAACTC |
| --- | --- |
| NOLC1-R | CCTGTAACTTTCGCTCTGGGA |
| CDKN1A-F | TGTCCGTCAGAACCCATGC |
| CDKN1A-R | AAAGTCGAAGTTCCATCGCTC |
| BAX-F | CCCGAGAGGTCTTTTTCCGAG |
| BAX-R | CCAGCCCATGATGGTTCTGAT |
| PTEN-F | AGGGACGAACTGGTGTAATGA |
| PTEN-R | CTGGTCCTTACTTCCCCATAGAA |
| FAS-F | AGATTGTGTGATGAAGGACATGG |
| FAS-R | TGTTGCTGGTGAGTGTGCATT |

**siRNA**

| si-NC | UUCUCCGAACGUGUCACGU |
| --- | --- |
| si-h-NOLC1-1 | CAAGAAGACUGUACCUAAA |
| si-h-NOLC1-2 | CCAAGAAUUCUUCAAAUAA |
| si-h-NOLC1-3 | CAUCUAAGUCUGCAGUUAA |
| si-h-P53-1 | GCAUCUUAUCCGAGUGGAAGGTT |
| si-h-p53-2 | ACUACAACUACAUGUGUAACATT |
| si-h-p53-3 | AGCGAGCACUGCCCAACAACATT |

**Cisplatin resistant cell line construction**

MGC-803 cells were treated with cisplatin (1 μM) for 3 month, then treated cells with cisplatin (5 μM) for 3month.

**Cell transfection and lentiviral infection**

MKN-45 and MGC-803 cells with stable NOLC1 expression at lower levels were generated by transducing a lentiviral vector followed by the ORF of NOLC1. Selected stable cells with puromycin (2 μg/ml) for 48 hours.

**Molecular docking**

Downloaded the crystal structure of P53 (PDB code: 8E7B) from the RCSB Data Bank (<http://www.pdb.org>). The NOLC1 protein structure was constructed by HDOCK based on its amino acid sequence. Using HDOCK software, each protein was set to be rigid, the docking contact sites were set to the full surface, and the resulting conformations were set to 100 after docking. The docking score was calculated based on the expert iterative scoring function ITScorePP. A more negative docking score implies a more likely binding model. In this study, the most negative energy conformation was selected by the evaluation function and optimized by the minimization module in the MOE 2019.1 software platform to solve the unreasonable contacts in the spatial structure that may occur in rigid docking. Amber10:ETH was selected as the force field for energy minimization, and water molecules were selected for the solvation model. The optimization method was divided into two steps: steepest descent and conjugate gradient, and the maximum number of iterations was 5000. Pymol2.1 software was used to visualize and analyze the model.
